# Selective PPAR-α activation with pemafibrate attenuates macrophage-mediated progression of calcific aortic valve disease

**DOI:** 10.64898/2025.12.17.694919

**Authors:** Mandy E. Turner, Yuto Nakamura, Takeshi Tanaka, Mark C. Blaser, Taku Kasai, Adrien Lupieri, Shinsuke Itoh, Rile Ge, Katelyn A. Perez, Rei Itagawa, Luisa Weiss, Takehito Okui, Yusuke Sasaki, Aruna D. Pradhan, Peter Libby, Paul M. Ridker, Sasha A. Singh, Masanori Aikawa, Elena Aikawa

**Affiliations:** Center for Interdisciplinary Cardiovascular Sciences, Cardiovascular Division, Department of Medicine, Brigham and Women’s Hospital, Harvard Medical School, Boston, MA, USA; Center for Excellence in Vascular Biology, Division of Cardiovascular Medicine, Department of Medicine, Brigham and Women’s Hospital, Harvard Medical School, Boston, MA, USA; Channing Division of Network Medicine, Department of Medicine, Brigham Women’s Hospital, Harvard Medical School, Boston, MA, USA; Center for Cardiovascular Disease Prevention, Division of Preventive Medicine, Department of Medicine, Brigham and Women’s Hospital, Harvard Medical School, Boston, MA, USA; Division of Cardiovascular Medicine, Department of Medicine, Brigham and Women’s Hospital, Harvard Medical School, Boston, MA, USA

**Author notes:** Corresponding author: Elena Aikawa, MD, PhD; Center for Life Sciences Boston Bldg., 17th Floor, 3 Blackfan Street, Boston, MA 02115. Turner ME and Nakamura Y contributed equally to this work.

**Keywords:** Aortic stenosis, calcific aortic valve disease, fibrate, pemafibrate, PPARα, macrophages, aortic valve wire injury model

## Abstract

**BACKGROUND:** Calcific aortic valve disease (CAVD) compromises valve compliance and cardiac hemodynamics leading to aortic stenosis (AS) and cardiovascular dysfunction. With treatment for severe AS limited to valve replacement and no effective pharmacotherapies, new interventions are urgently needed for patients. This study evaluated pemafibrate, a selective peroxisome proliferator-activated receptor alpha (PPARα) activator as a novel therapeutic for CAVD and AS.

**METHODS AND RESULTS:** In an aortic valve wire injury (AVWI) model of AS in Ldlr^⁻/⁻^ mice, pemafibrate administration (0.2 mg/kg/day) for 15 weeks improved aortic valve function and reduced valvular calcification by 39% (p<0.001), accompanied by reduced leaflet inflammation and CD68⁺ macrophage infiltration. These effects were independent of changes in plasma triglyceride levels. *In vitro*, pemafibrate suppressed inflammation-mediated calcification of primary human valvular interstitial cells (VICs) by modulating macrophage-derived secreted factors, identifying macrophage–VIC crosstalk as a key disease mechanism. Direct treatment of macrophages with pemafibrate, or exposure to serum from pemafibrate-treated participants in the PROMINENT randomized controlled trial, shifted macrophages toward a less inflammatory and less chemotactic phenotype. Proteomic analyses of patient serum substantiated these findings by reflecting a systemic reduction in inflammatory parameters and monocyte activation. Network integration of the *in vitro* derived pemafibrate-responsive proteome with human calcified AV tissue proteomes identified aberrant protein translation (GNB2L1, GSPT1) and disrupted bioenergetics (MYDGF, PDIA4) as potential clinically relevant pemafibrate-responsive pathways and effector proteins relevant to AS progression.

**CONCLUSIONS:** Pemafibrate slows experimental AS progression and valve calcification through modulation of macrophage-VIC crosstalk, independent of lipid lowering. These findings support further evaluation of pemafibrate as a potential pharmacological approach for CAVD and support further testing in randomized clinical trials.

## Introduction

Calcific aortic valve disease (CAVD) affects more than 25% of individuals over the age of 65^1^. CAVD can progress to aortic stenosis (AS), whereby calcification of the aortic valve (AV) impairs leaflet motion and obstructs left ventricular outflow leading to severe cardiac dysfunction and heart failure^2^. Without intervention, severe AS carries a high mortality rate, with only half of patients surviving beyond two-years^3^. Invasive and costly AV replacement surgery and transcatheter AV implantation are the sole treatment options for patients. Further, traditional cardiovascular pharmacotherapies such as statins have proven ineffective for the prevention of AS^4^. Taken together, an urgent need exists for medical options to prevent or treat CAVD in patients^5^.

CAVD pathogenesis is multifactorial and arises, in part, from a complex pro-inflammatory local environment characterized by valvular cell differentiation, immune cell infiltration, and lipid deposition^4,6^. Valvular interstitial cells (VICs) undergo myofibroblastic and osteogenic differentiation which promote leaflet fibrosis and calcification^7^. Factors secreted from local cells and cell-cell communication are central to calcification promotion within the AV^8–10^. Secreted extracellular vesicles from both osteogenic VICs and macrophages can initiate calcification in the extracellular matrix (ECM)^10–13^. Additionally, accumulating evidence suggests that macrophages can initiate adaptive immune responses to expedite calcification progression through intercellular communication with VICs^14–17^. Macrophages secrete pro-inflammatory cytokines such as interferon-gamma (IFNγ), tumor necrosis factor-alpha (TNF-α), and interleukin-6 (IL-6) that have been associated with calcification progression in VICs^14,15,18^. Exploring mechanisms to attenuate pro-inflammatory responses within the AV could therefore provide insight into effective approaches for preventing CAVD.

The fibrate drug class lowers circulating triglycerides (TGs) through activation of hepatic peroxisome proliferator activated receptor α (PPARα) to increase lipoprotein lipase^19^. Pemafibrate is highly selective for PPARα and, in addition to lipid lowering, we have reported that pemafibrate reduces pro-inflammatory properties of macrophages and attenuates vein graft disease, in-stent stenosis, and metabolic dysfunction-associated steatohepatitis independently of its TG-lowering action^20–23^. The Pemafibrate to Reduce Cardiovascular Outcomes by Reducing Triglycerides in Patients with Diabetes (PROMINENT) randomized controlled trial demonstrated the safety of pemafibrate among patients with type 2 diabetes and mild-to-moderate hypertriglyceridemia^24^. Fibrates have yet to be investigated as a therapy for CAVD. Given the recognized role of inflammation in driving CAVD pathogenesis, we hypothesized that pemafibrate may attenuate valvular calcification through its anti-inflammatory effects.

Using *in vivo* and *in vitro* models alongside translational integration of pemafibrate-treated clinical trial participants and calcified human AV tissue proteomic profiling, this study is the first to demonstrate that PPARα activation with pemafibrate inhibits valvular calcification. We identified a macrophage-VIC crosstalk axis as the primary mechanism. These findings position pemafibrate as a potentially effective therapeutic agent for the treatment of AS that merit future clinical investigation.

## Results

### Pemafibrate slowed progression of aortic stenosis in an AVWI mouse model

The effect of the PPARα-selective activator, pemafibrate, was assessed in an established model of AS characterized by local inflammation using AV wire injury (AVWI) in low-density lipoprotein receptor-deficient (Ldlr^-/-^) mice^7,25^. Mice were maintained with or without a clinically relevant dose of pemafibrate (0.2 mg/kg/day)^20^ on either a normal diet (ND) or high-fat diet (HFD) for 15 weeks (**Figure 1A**). Body weight was reduced in pemafibrate-treated mice compared to controls without concomitant reductions in food intake in both ND and HFD conditions (**Figure 1B-C**). These results are consistent with previous reports that pemafibrate suppressed HFD-induced body weight gain in mice^26,27^. Pemafibrate did not affect heart weight or heart weight-to-body weight ratio at endpoint (Week 15; **Supplementary Figure S1**). As expected, pemafibrate suppressed circulating TG levels without changing the total cholesterol or phosphate levels in both the ND and HFD groups (**Figures 1D-E**, and **Supplementary Figure S2A**). Pemafibrate decreased the total plasma calcium levels in HFD mice (**Supplementary Figure S2B**).

**Figure 1:**
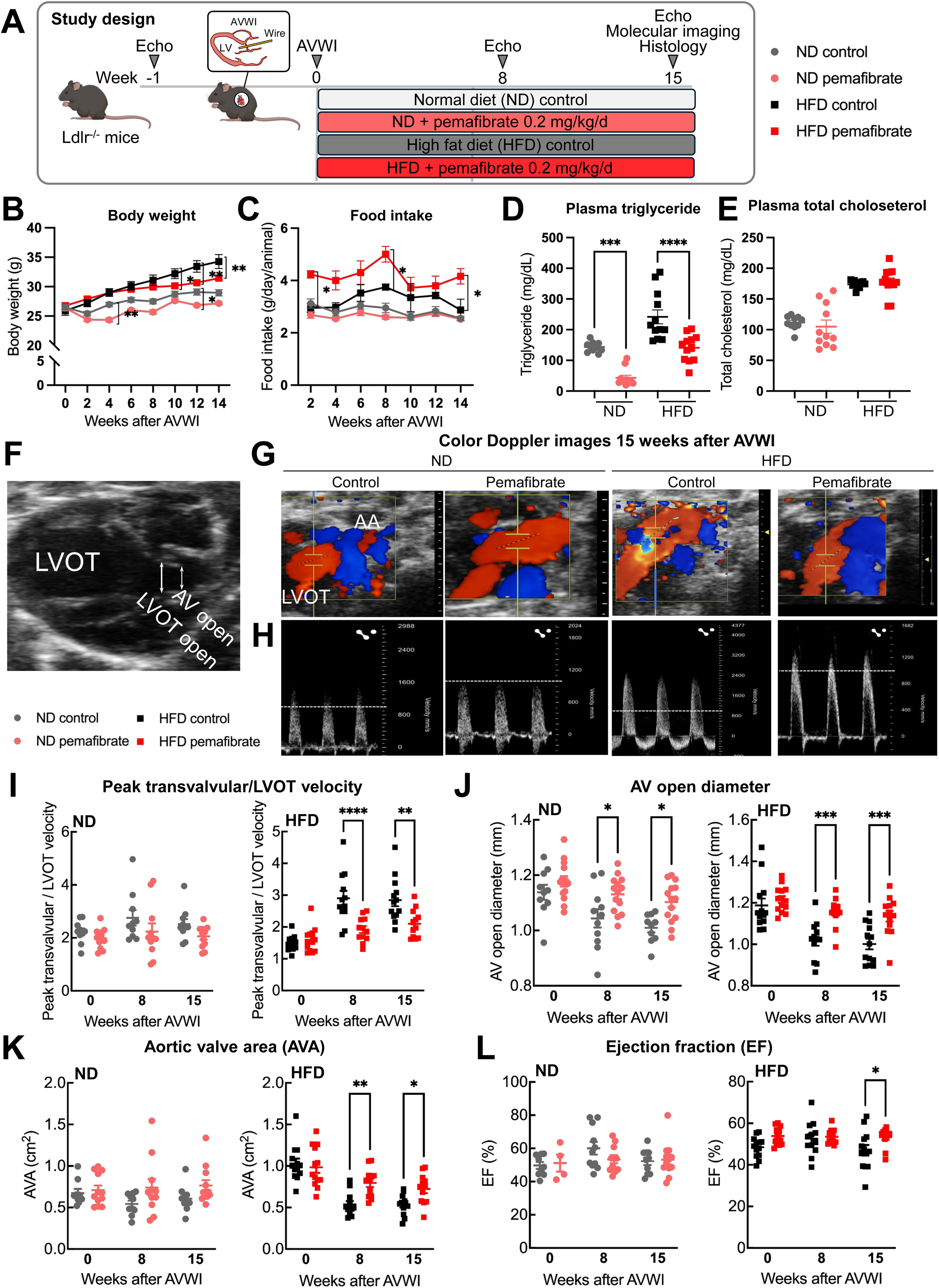
Pemafibrate improved aortic stenosis in an aortic valve wire injury mouse model. **(A)** *In vivo* study design: Ldlr-/- mice underwent AVWI surgery and were randomly assigned to a normal diet (ND) or a high-fat diet (HFD) group, with or without pemafibrate (0.2 mg/kg/day). **(B)** Body weight. **(C)** Food intake. **(D-E)** Quantification of plasma triglyceride and total cholesterol at 15 weeks. **(F-H)** Representative images of aortic valve (AV) area (F), color doppler (G), and transvalvular peak velocity (H). The white dotted line represents a velocity of 1,000 mm/s. **(I-L)** Echocardiography measurement of the ratio of transvalvular blood velocity/left ventricular outflow tract (LVOT) velocity (I), AV open diameter (J), aortic valve area (K), and ejection fraction (I). Mean±SEM. (B)-(E) n=12/group, (I)-(L) n=4-12/group. Ordinary two-way ANOVA followed by Bonferroni *post hoc* test performed for statistical analysis between control and pemafibrate in ND and HFD. *p<0.05, **p<0.01, and ***p<0.001.

The impact of pemafibrate on cardiac and valvular function was assessed via echocardiography before wire injury as well as at 8- and 15-weeks post-wire injury (**Figure 1F-H Supplementary Table S1**). AS and impaired valve function were induced following AVWI in all groups, with significant elevations of peak transvalvular velocity normalized to left ventricular outflow tract (LVOT) velocity and reduced AV open diameter. These functional deficits were attenuated by pemafibrate treatment at both 8 and 15 weeks (**Figure 1I-J**). With HFD, pemafibrate also improved AV area (AVA, **Figure 1K**). Pemafibrate increased the ejection fraction (EF) compared to control only in the HFD group at 15 weeks post-AVWI (**Figure 1L**). Overall, pemafibrate treatment consistently attenuated AS-associated echocardiography parameters in the HFD group.

### Pemafibrate attenuated valvular calcification and inflammation in AVWI mice fed HFD

*In vivo* near-infrared fluorescence (NIRF) molecular imaging demonstrated that pemafibrate significantly attenuated AVWI-induced calcification and matrix metalloprotease (MMP) activity, suggestive of ECM remodeling, in the AVs of HFD fed mice (**Figure 2A-D**). Pemafibrate also reduced valve leaflet thickness and valvular CD68^+^ macrophage content, compared to untreated mice in both ND and HFD conditions (**Figure 2E-H**). In the AV, the presence of two inflammatory parameters, IL-12 which associates with cardiovascular calcification^13–16^ and S100A9, which contributes to inflammation-associated microcalcification formation within atherosclerotic plaques^28,29^, were higher in HFD AVWI mice, and pemafibrate treatment attenuated these proinflammatory features (**Figure 2I-L**). Consistent with the *in vivo* NIRF imaging data, high-resolution histopathological imaging of AVs demonstrated punctate areas of calcification in HFD AVWI mice, which were attenuated by pemafibrate treatment compared to untreated mice (**Supplementary Figure S3**). These results demonstrate that pemafibrate ameliorated AS AVWI-induced morphological and local inflammatory changes in the AV.

**Figure 2:**
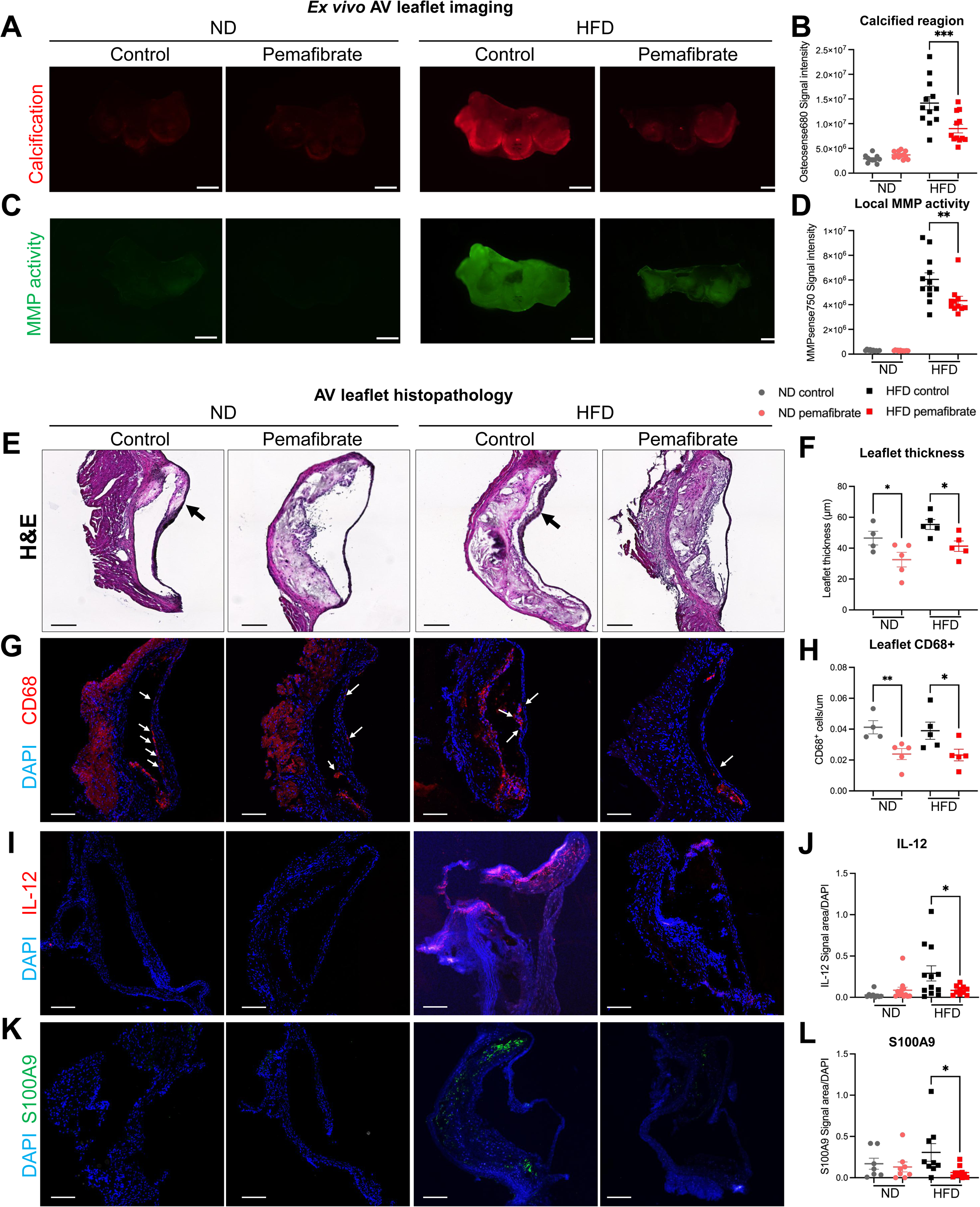
Pemafibrate suppressed calcification and pro-inflammatory changes of the aortic valve. **(A)** Representative dissected aortic root (3-leaflet view) stained with Osteosense680. Scale bar=1 mm. **(B)** Calcified region quantification in the whole valve tissue (Osteosense680 signal intensity). **(C)** Representative local MMP activity stained with MMPsense750. Scale bar=1 mm. **(D)** Local MMP activity quantification in the whole valve tissue (MMPsense750 signal intensity). **(E)** Representative images of hematoxylin & eosin (H&E) staining. Red asterisk indicates calcified region. The black arrow indicates thickening of the valve leaflet. **(F)** Quantification of AV leaflet thickness. **(G)** Representative images of CD68 immunostaining (Blue, DAPI: Red, CD68). The white arrows indicate CD68^+^ cell infiltration sites. **(H)** Quantification of CD68^+^ cell infiltration in AV leaflets. Data was normalized by the length of AV leaflet. **(I)** Representative images of IL-12 immunostaining (Blue, DAPI; Red, IL-12). **(J)** IL-12 signal area normalized by DAPI in the whole AV section. **(K)** Representative images of S100A9 immunostaining (Blue, DAPI; Green, S100A9). **(L)** S100A9 signal area normalized by DAPI in the whole AV section. Mean±SEM. (B) and (D) n=10-12. Scale bar=1 mm (F) and (H) n=4-5/group. (H), (J), and (L) n=10-12/group. Scale bar=200 μm. Ordinary two-way ANOVA followed by Bonferroni *post hoc* test performed for statistical analysis between control and pemafibrate-treated in ND and HFD. *p<0.05, **p<0.01, ***p<0.001, and ****p<0.0001.

### *In vivo* effects of pemafibrate on AS did not correlate with TG levels

Pemafibrate is a potent TG-lowering drug. The various changes in AV with pemafibrate administration, including peak LVOT velocity, AV open diameter, and AVA, however, did not correlate with plasma TG levels (**Supplementary Figure S4**). Histopathological parameters of AS and inflammation also did not correlate with plasma TG levels (**Supplementary Figure S5**). These *in vivo* results suggest that pemafibrate suppressed AS phenotype and local AV inflammation, at least in part, independently of its TG-lowering effects, motivating *in vitro* mechanistic assessment of pemafibrate on local cells that contribute to CAVD pathogenesis.

### Pemafibrate treatment did not directly attenuate valvular interstitial cell calcification *in vitro*

Given that pemafibrate inhibited AV calcification in the AVWI model (**Figures 1-2**), we hypothesized that pemafibrate may directly suppress calcification of primary VICs isolated from AS patient valves. However, pemafibrate administered at 0.1-10 μM failed to reduce osteogenic media-induced calcification as measured by alizarin red staining in VICs (n=6 donors), indicating that the anti-calcification effects observed *in vivo* are mediated through intermediate cellular mechanisms rather than direct action on VICs (**Supplementary Figure S6**).

### Pemafibrate suppressed VIC calcification through macrophage-secreted factors

Given that pemafibrate treatment reduced local inflammation and macrophage accumulation in the AVWI model, we investigated a potential role for macrophage-VIC communication in mediating valvular calcification (**Figure 3A**). To address this, we assessed the effects of a macrophage secretome by applying conditioned media (CM) from THP-1 macrophage-like cells pre-treated with pemafibrate (0.1 μM, 1 μM, 10 μM) or DMSO to human VICs (n=6 donors, **Figure 3A**).

**Figure 3:**
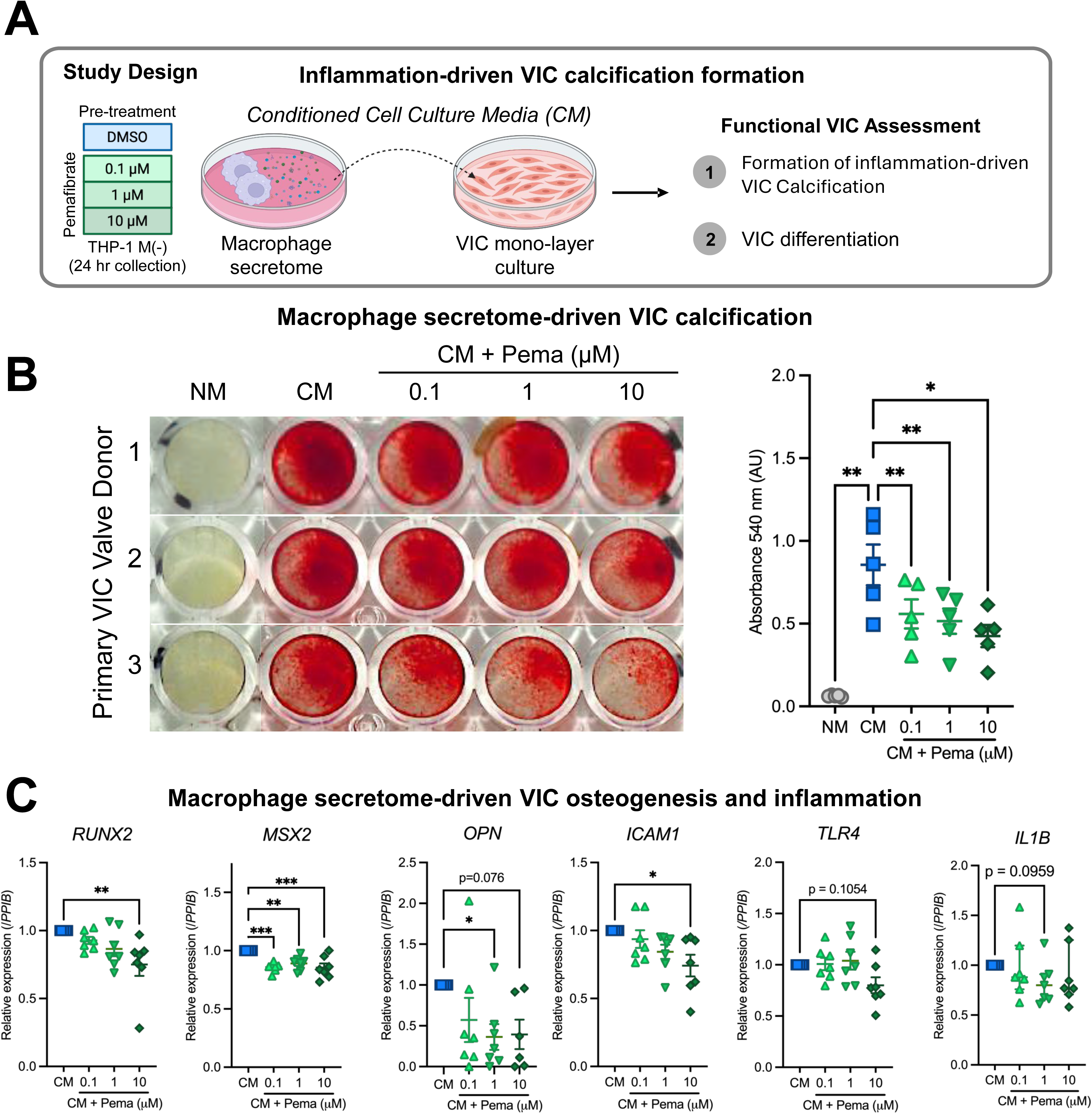
Pemafibrate suppressed calcification of human primary valvular interstitial cells through pleiotropic effects of macrophages. (**A**) Schematic of *in vitro* assay of primary VICs exposed to conditioned media (CM) from non-polarized THP-1 macrophage-like cells, M (-), pre-treated with 0, 0.1, 1, or 10 μM pemafibrate (Pema). (**B**) Alizarin red staining and quantification for calcium deposition in VICs exposed to CM with or without pemafibrate pre-treatment or non-cell exposed control media (NM) for 28-35 days. Mean ± SEM. N=5 donors. Repeated measures one-way ANOVA with Dunnett’s multiple comparison test. *p<0.05, **p<0.01. (**C**) Osteogenic (*RUNX2*, *MSX2*, *OPN*) and inflammatory (*ICAM1*, *TLR4*, *IL1B*) gene expression after 7 days of exposure to CM. One-way ANOVA with *post hoc* Dunnett’s multiple comparison test. *p<0.05, **p<0.01, ***p<0.001.

Exposure to the macrophage secretome, without additional osteogenic stimulus, induced calcium deposition as indicated by positive alizarin red staining within all donors and increased alkaline phosphatase (ALP) activity (**Figure 3B** and **Supplementary Figure S7**). Pre-treatment of THP-1 macrophage-like cells with pemafibrate dose-dependently attenuated calcification (**Figure 3B**). Genes characteristic of osteoblastic differentiation (*MSX2, RUNX2, OPN*) and VIC inflammation pathway activation (*TLR4, ICAM1, and IL1B*) decreased in VICs when the CM was taken from THP-1 macrophage-like cells pre-treated with pemafibrate (**Figure 3C**). This demonstrates that pemafibrate attenuates both the osteogenic and inflammatory VIC phenotypes via macrophages, processes previously linked to AV calcification^14^.

### Pemafibrate pre-treatment of macrophages rescued proteomic features associated with extracellular matrix remodeling, cellular metabolism, and protein translation in VICs

It is noteworthy that the macrophage secretome alone exerted a robust pro-calcific influence without any supplementary stimuli, yet the specific mechanisms driving this inflammatory-mediated osteogenic response remain undefined. We used mass spectrometry proteomics to characterize the molecular response of VICs to macrophage-derived CM, and to assess which of these features were attenuated by pemafibrate. This enabled us to pinpoint specific pathways through which macrophage-VIC crosstalk promotes calcification (**Figure 4A**). Principle component analysis (PCA, **Figure 4B**) demonstrated that after 7 days there was a distinct effect of CM on the VIC proteome compared to NM, with a further effect of macrophage pre-treatment with pemafibrate (CM + 0.1-10 μM pemafibrate).

**Figure 4:**
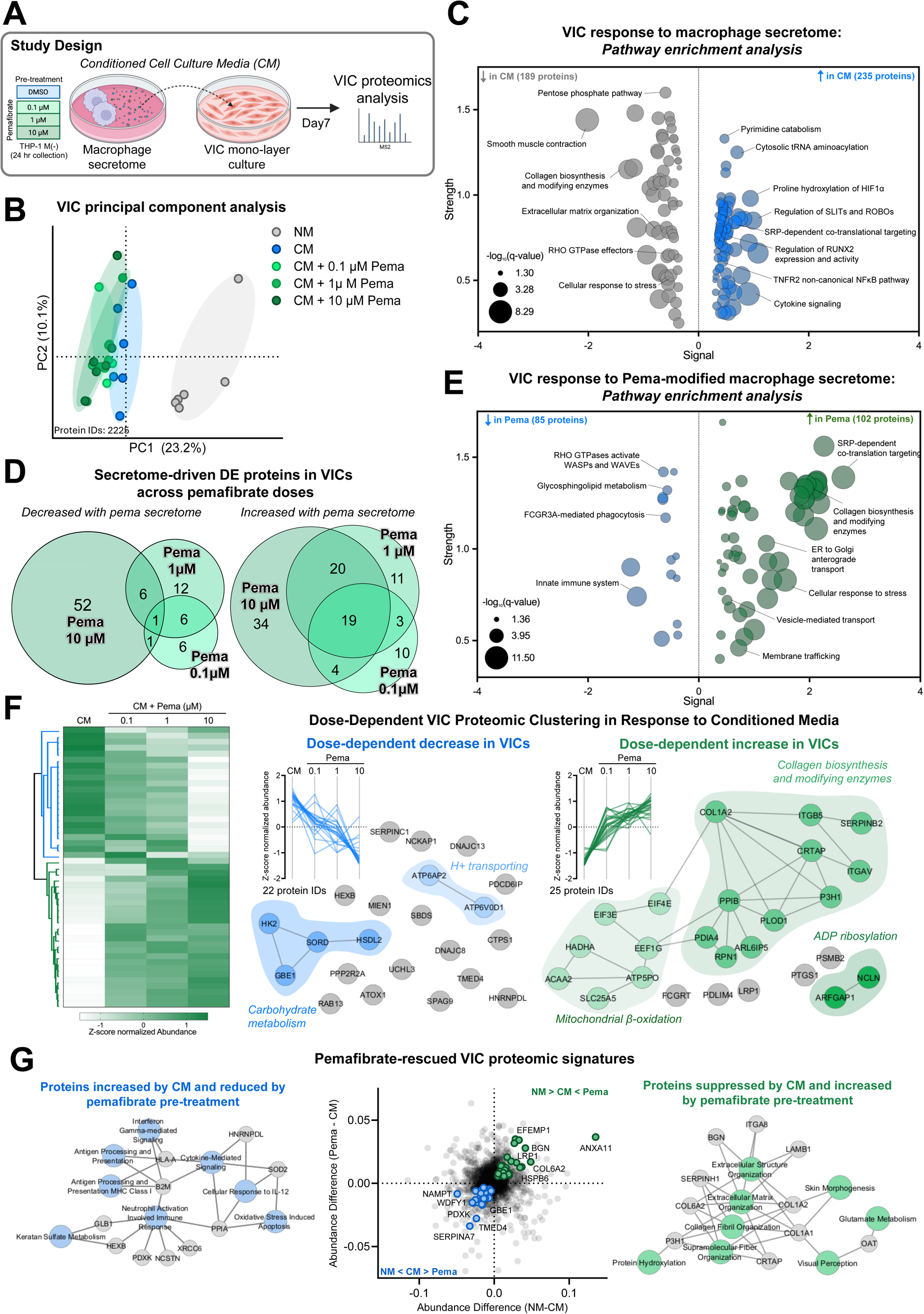
Valvular interstitial cells exposed to the secretome of macrophages underwent an inflammatory and osteogenic proteomic shift, which was attenuated by pre-treating the macrophages with pemafibrate. **(A)** Schematic of *in vitro* assay of primary VICs (N=6 donors) exposed to conditioned media (CM) from non-polarized THP-1 macrophage-like cells, M (-), pre-treated with 0, 0.1, 1, or 10 μM pemafibrate (Pema). **(B)** VIC principal component (PC) analysis. **(C)** VIC cellular response to macrophage secretome. Reactome pathway enrichment analysis of proteins differentially enriched in CM compared to control NM conditions with q<0.05 and StringDB-generated scores. ANOVA followed by *post hoc* Tukey HSD test to determine differentially enriched proteins between each group (q<0.05). Input gene ID list (Supplementary Table S3) and all Reactome pathways and annotations (Supplementary Table S4). **(D)** Dose-wise overlap of VIC proteins either increased or decreased by macrophage pemafibrate pre-treatment compared to CM. ANOVA followed by *post hoc* Tukey HSD test to determine differentially enriched proteins between each group (q<0.05, gene IDs Supplementary Table S5) and **(E)** Reactome pathway enrichment analysis of any differentially enriched protein with q<0.05 and StringDB-generated scores. Input gene ID list (Supplementary Table S6) and all Reactome pathways and annotations (Supplementary Table S7). **(F)** Hierarchical clustering of dose-responsive proteins. Hierarchical clustering ANOVA-significant (q<0.05) proteins of conditioned media +/-pemafibrate. Dose-dependent decreasing proteins (22) and increasing proteins (25) StringDB protein-protein interaction network with k-means clustering and annotation (Supplementary Table S8). **(G)** Identification of pemafibrate-responsive proteins that return to a control state. Magnitude and direction of differential enrichment between NM and CM plotted against CM and 10μM pemafibrate. Proteins with significantly increased abundance in response to CM and subsequently significantly decreased with pemafibrate macrophage-pre-treatment are highlighted in blue, and proteins with significantly reduced abundance in response to CM and subsequently significantly increased abundance with pemafibrate macrophage-pre-treatment are highlighted in green (p<0.05, Supplementary Table S9). Top 8 GO Biological Processes enrichment analysis and network using EnrichrKG. Unabridged pathway annotations in Supplementary Table S9.

Consistent with the pro-calcific effect of the macrophage secretome on VICs (**Figure 3**), CM induced inflammatory and pro-osteogenic responses in the VIC proteome, as well as perturbations in proteins associated with endoplasmic reticulum protein translation (**Figure 4C**). Proteins associated with the innate immune response and cytokine signaling (HLA-A, B2M, HMGB1, proteosome subunits: PSMB8/LMP7, PSME3) increased. Additionally, calcification-associated proteins, RUNX2, a master transcriptional regulator of osteogenesis, and TNFRSF11B (Osteoprotegerin, OPG), a negative regulator of osteoclastogenesis, were systemically elevated in individuals with cardiovascular calcification^30^ (q<0.05, **Supplementary Table S3-S4**). PTGS1/S2 (COX-1, COX-2), which has been causally implicated in inflammatory responses and valvular calcification and Rho-GTPase-RhoROCK signalling^31^, was also increased in response to CM.

We next defined the modulatory effect of pemafibrate pre-treatment on macrophage secretome-induced VIC phenotypic changes. The highest doses of pemafibrate pre-treatment of macrophages induced the most robust VIC proteomic shifts compared to CM (q<0.05, **Figure 4D and Supplementary Table S5**). Compared to CM alone, addition of pemafibrate decreased VIC proteins associated with innate immune response. These included several proteins implicated in calcification, such as MAPK (ERK) and PKM (**Supplementary Table S6-S7**)^32,33^. Pathways associated with Rho GTPase signaling, which were increased in CM- vs NM-treated VICs, were normalized by pre-treatment of macrophages with pemafibrate. Further, there was a reduction in several vesicle-trafficking related proteins, a process known to be central to calcification (RANGAP1, RBM8A, TMED4, TRIM16, AP2A2, ARHGAP17, DNAJB4, GM2A, PSAP, ANXA3) as well as S100-family calcium binding proteins (S100A4, S100A10)^8,10,12^.

Using hierarchical clustering, we identified two VIC protein clusters which dose-dependently decreased (n=22, q<0.05) or increased (n=25, q<0.05, **Figure 4F**). Proteins that dose-dependently decreased were associated with metabolic programming and glycolytic shifts (carbohydrate metabolism: HK2, GBE1) and vacuolar proton-transporting V-type ATPase, which have been implicated in pro-calcific Wnt signaling and lysosomal acidification^34,35^ (**Supplementary Table S8**). Pemafibrate pre-treatment of the THP-1 macrophage-like cells dose-dependently increased proteins associated with the ECM, specifically collagen maintenance in VICs. Collagen fragmentation and disorganization is a central feature of ectopic calcification^36,37^.

We defined which specific proteomic CM-induced shifts of the VICs are rescued when the THP-1 macrophage-like cells were pre-treated with pemafibrate (**Figure 4G** and **Supplementary Table S9**). Pemafibrate pre-treatment attenuated the elevation of VIC proteins associated with inflammation and cytokine signaling (HLA-A, PPIA, B2M), and restored proteins related to ECM homeostasis and stability, including various collagen sub-types and binding proteins (COL1A1, COL1A2, COL6A2, BGN), and proteins involved in collagen folding and post-translational modifications (FKBP10, FKBP9, CRTAP, P3H1, P3H3, SERPINH1). BGN deficiency has been associated with valvular calcification attenuation^38,39^.

Importantly, only a small fraction of THP-1-derived proteins detected in CM overlapped with proteins identified in the VIC differential enrichment analysis (n=23/223, 10.3%; **Supplementary Figure S8, Supplementary Table S10**), suggesting that the VIC proteomic shifts do not originate from macrophage CM. These overlapping proteins were primarily associated with protein binding, innate immune function, and ECM interactions (**Supplementary Table S11-S12**).

### Pemafibrate and serum from patients treated with pemafibrate shifted macrophage phenotype to a less inflammatory and migratory state

Given that pemafibrate-pretreatment of THP-1 macrophage-like cells attenuated the calcific potential of the CM in VICs, we assessed the phenotype of THP-1 cells after pemafibrate treatment under pro-inflammatory conditions (IFNγ+LPS) using targeted and discovery-based approaches (**Figure 5A**). Pemafibrate induced a robust shift in the activated THP-1 cellular proteome (n=366/2551 differentially enriched proteins, q<0.05, **Figure 5B-C, Supplementary Table S13).** Pemafibrate reduced proteins associated with pro-inflammatory pathways such as interferon signaling (STAT1), as well as chemokine signaling (MAPK, HCK, BST2, **Figure 5D, Supplementary Table S14-S16)**. Proteins increased by pemafibrate, a highly selective PPARα activator, were most associated with an increase in PPARα-activated gene expression pathways as detected by functional enrichment analysis of the entire proteome. Given the modulation of inflammatory and chemotaxis-related pathways, we performed targeted gene expression assessment of canonical markers, and confirmed consistent reduction across both pathways, including marked reductions in *S100A9, IL-12,* and matrix remodeling enzymes *MMP2/9,* consistent with the decrease of those proteins in valve leaflets of pemafibrate-treated AVWI mice **(Figure 5E)**.

**Figure 5:**
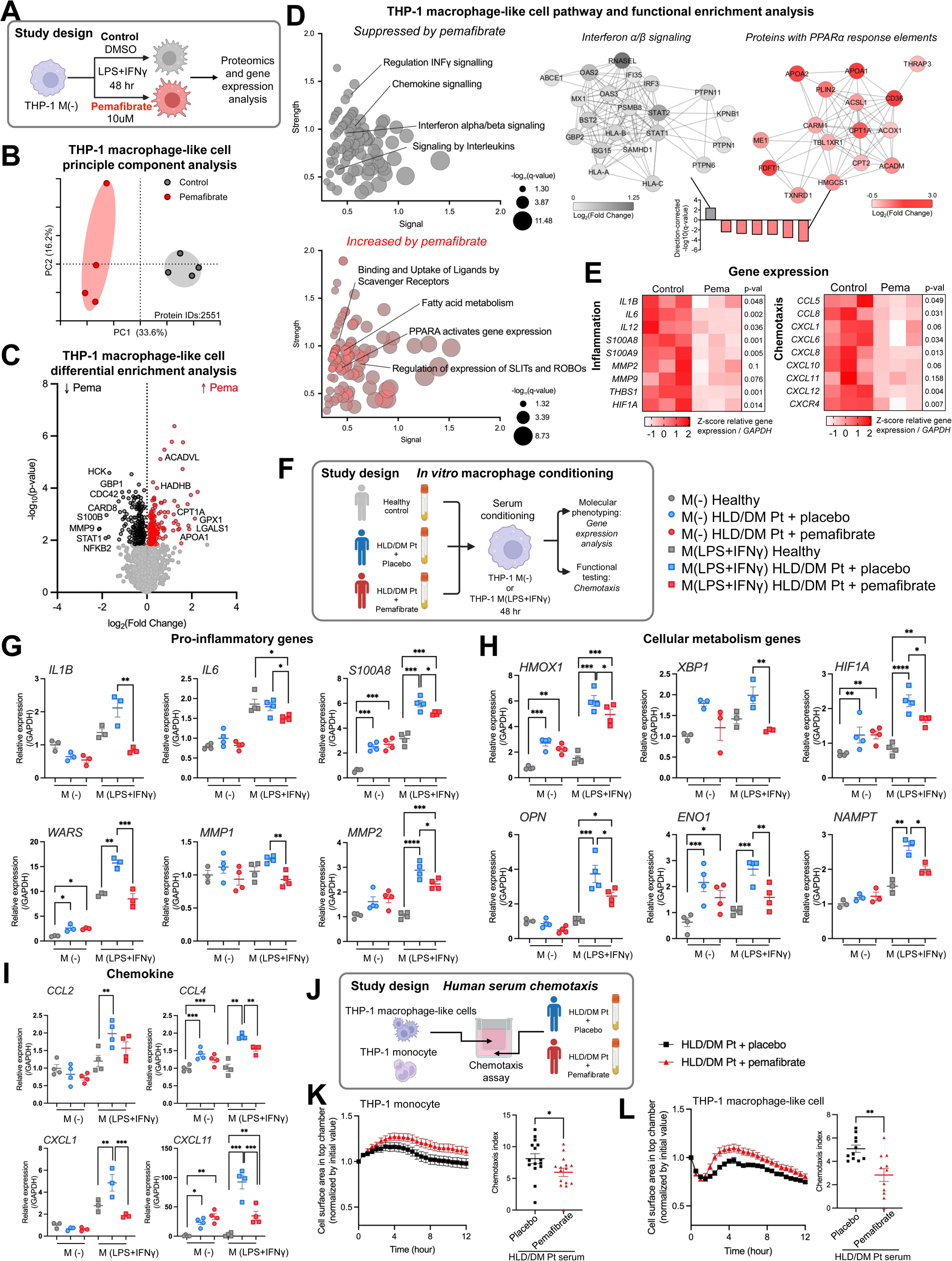
Pemafibrate and serum from patients treated with pemafibrate modulated macrophage phenotype to a less inflammatory and migratory state. **(A)** Schematic of *in vitro* assay of THP-1 macrophage-like cells stimulated to a pro-inflammatory phenotype with LPS and IFNγ (M(LPS+IFNγ), N=4) exposed to vehicle DMSO (control) or 10 μM pemafibrate. **(B)** THP-1 principal component (PC) analysis. **(C)** THP-1 cellular response to pemafibrate. Volcano plot of differential enrichment analysis using T-test. Highlighted in red and black are proteins with FDR q<0.05 (Supplementary Table S13). **(D)** Pathway and functional enrichment analysis of THP-1 pemafibrate response. Reactome pathway enrichment analysis of proteins differentially enriched in controls or pemafibrate with q<0.05 and StringDB-generated scores. Input gene ID list (Supplementary Table S14) and all Reactome pathways and annotations (Supplementary Table S15). Functional enrichment analysis using fold change ranking in StringDB identifies 7 enriched pathways (FDR q<0.05). Protein-protein interaction networks of the most significantly suppressed pathway (interferon α/β signaling) and the most enriched pathway by pemafibrate (proteins with PPARα genetic response elements), with node colors corresponding to fold change. All Reactome pathways and annotations (Supplementary Table S16). **(E)** Gene expression analysis of canonical inflammation- and chemotaxis-associated genes (n=3). Gene expression plotted as individual replicates of Z-score normalized gene expression relative to GAPDH, and p-value calculated using T-test. **(F)** Schematic of *in vitro* assay of non-polarized THP-1 macrophage-like cells, M(-), or M(LPS+IFNγ) stimulated with HLD/DM patient-derived serum. **(G)-(I)** Gene expression analysis in M(-) or M(LPS+IFNγ) exposed to healthy control or HLD/DM patient-derived serum (n=3-4). Pro-inflammatory genes (G), cellular metabolism genes (H), and chemokines (I) were measured. **(J)** Schematic of chemotaxis assay using HLD/DM patient-derived serum. **(K)-(L)** In either THP-1 macrophage-like cells (K) or monocytes (L), the cell surface area in top chamber was normalized by the initial cell area, and plotted in 12-hour time window (n=11-15). The chemotaxis index was calculated by the area under the curve value of each group. Mean±SEM. Ordinary two-way ANOVA followed by Bonferroni *post hoc* test performed for statistical analysis.

To evaluate translational potential of our pre-clinical evidence, we used serum from patients with hyperlipidemia and type 2 diabetes (HLD/DM Pt) treated with pemafibrate or placebo for 4 months in the PROMINENT clinical trial^24^ (pooled N=12 donors/group) alongside commercial serum from healthy donors, to stimulate THP-1 macrophage-like cells under both non-polarized M(-) and pro-inflammatory M(LPS+IFNγ) conditions **(Figure 5F)**. Targeted assessment of inflammatory- (*IL1B, S100A8, WARS, MMP1, MMP2*), cellular metabolism- (*HMOX1, XBP1, HIF1A, OPN, ENO1, NAMPT),* and chemotaxis- (*CCL2, CCL4, CXCL1, CXCL11*) related macrophage gene expression demonstrated a consistent increase in response to serum from HLD/DM Pts compared to healthy controls, followed by a marked attenuation with pemafibrate treatment. These effects were more pronounced in the already pro-inflammatory M(LPS+IFNγ) cells **(Figure 5G-I)**.

Since pemafibrate-treated AVWI mice had reduced CD68+ macrophage content in the AV leaflets and consistent identification of chemotaxis-related genes and proteins in response to pemafibrate, we assessed the functional impact of serum derived from pemafibrate-treated HLD/DM Pts on macrophage chemotaxis **(Figure 5J)**. Because pemafibrate was administered twice daily in the PROMINENT trial, we assessed chemotaxis over a 12-hour period. Pemafibrate treatment of patients with HLD/DM significantly reduced chemotaxis and cell migration, compared to placebo, when the serum was used as the attractant in both THP-1-derived monocytes **(Figure 5K)** and macrophage-like cells **(Figure 5L)**.

### The serum proteome of pemafibrate-treated patients reflected attenuation of pro-inflammatory innate immune activation

To expand upon the *in vitro* evidence to further assess translational potential and mechanisms of pemafibrate in AS, we examined the serum proteome of HLD/DM Pt treated with pemafibrate or placebo for 4 months from PROMINENT (N=12/group, **Figure 6A)**^24^. Demographics of the 24 patients were comparable between treatment groups with respect to age, sex distribution, race, HbA1c %, and baseline lipid parameters **(Table 1)**. Placebo patients had a higher average BMI at study initiation (baseline) than the pemafibrate group. Following 4 months of pemafibrate treatment, as expected, TG and ApoC3 levels decreased in the pemafibrate-treated group whilst total cholesterol levels did not change.

**Figure 6:**
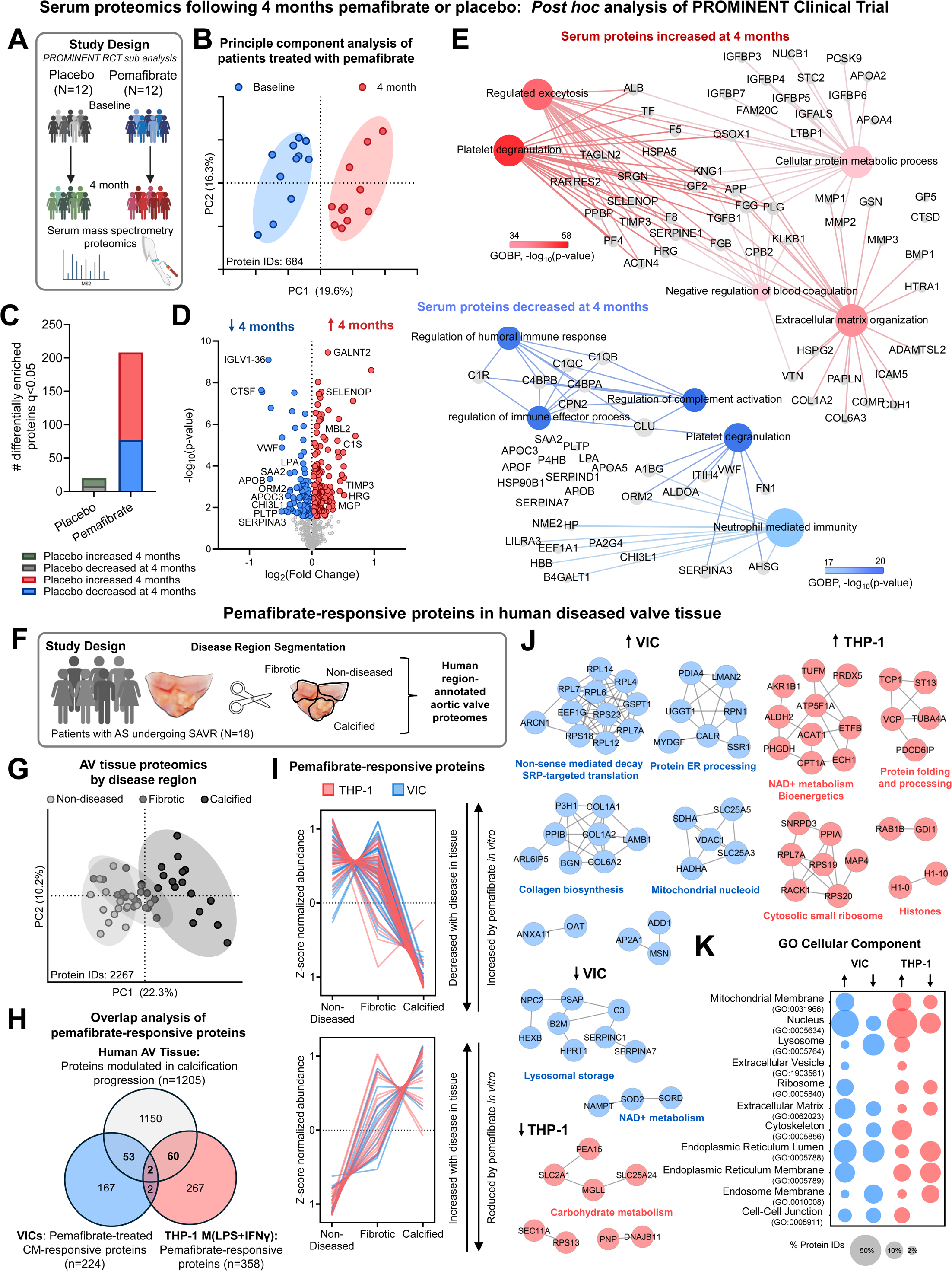
Four months of pemafibrate treatment was sufficient to induce a robust shift in the human serum proteome that reflects an attenuation of innate immune system activation and inflammation. **(A)** Study design of serum proteomics in HLD/DM patients before and after pemafibrate treatment. **(B)** Principal component analysis (PCA) of HLD/DM patients’ serum proteomics at baseline and post-pemafibrate treatment (4 months). **(C)** The number of differentially enriched proteins (q<0.05). **(D)** Volcano plot of differential enrichment analysis using T-test. Highlighted in red and black are proteins with FDR q<0.05 (Supplementary Table S17). **(E)** Gene Ontology Biological Processes (GOBP) protein network highlighting key proteins increased or decreased by pemafibrate (Supplementary Table S18). **(F)** Study design of disease region-segmented human AV tissue proteomics into non-diseased, fibrotic, and calcified regions from patients undergoing surgical AV replacement (N=18). **(G)** PCA of stage-segmented human AV tissue proteomes. **(H)** Overlap analysis of (i) differentially enriched proteins across disease stage of human AVs (N=18) ANOVA followed by *post hoc* Tukey HSD test to determine differentially enriched proteins between each group (q<0.05, Supplementary Table S21), (ii) differentially enriched proteins from THP-1 macrophage-like cells treated with pemafibrate or DMSO control (q<0.05, Supplementary Table S14 – Figure 3), and (iii) differentially enriched proteins in VIC cell lysates exposed to conditioned media from THP-1 macrophage like cells pre-treated with 0, 0.1, 1, or 10 μM pemafibrate (q<0.05, Supplementary Table S6 – Figure 5). Overlap signifies protein IDs that were increased across disease stage (C>ND, F>ND, C>ND) and then decreased by pemafibrate directly (THP-1) or indirectly (VIC) or decreased across disease stage (C<ND, F<ND, C<ND) and increased by pemafibrate directly (THP-1) or indirectly (VIC). **(I)** Profile plots of overlapping proteins from panel H separated by directionality. Full list of protein IDs in Supplementary Table S22. **(J)** Cell-type and directionality separated stringDB PPIs clustered using Markov cluster algorithm with a granularity parameter of 4. Singletons are not plotted. Select pathway enrichment of each cluster highlighted on figure. All annotations in Supplementary Table S23. **(K)** Bubble plot of Gene Ontology Cellular Compartment pathway identification of overlapping proteins via Enrichr. Size of bubble indicates the percentage of proteins identified in the ontology divided by the total number of proteins in the overlap analysis.

**Table 1:** Participants with hyperlipidemia and type 2 diabetes treated with pemafibrate or placebo for 4-months (N=24). Sub-analysis of PROMINENT randomized controlled trial.

|  |  | Placebo (n=12) | Pemafibrate (n=12) | p-value |
| --- | --- | --- | --- | --- |
| *Demographics and single measurement variables* |  |  |  |  |
| Age (yrs) |  | 66.6 ± 6.6 | 64.5 ± 10.3 | 0.56 |
| Sex (female) |  | 6 (50%) | 6 (50%) | 0.99 |
| Race | White | 9 (75%) | 8 (66.7%) | 0.88 |
|  | Multiracial | 1 (8.3%) | 1 (8.3%) |  |
|  | Other | 2 (16.7%) | 3 (25%) |  |
| Smoking | Current | 0 (0%) | 3 (25%) | 0.15 |
|  | Former | 7 (58.3%) | 4 (33.3%) |  |
|  | Never | 5 (41.7%) | 5 (41.7%) |  |
| Statin Use (yes) |  | 12 (100%) | 12 (100%) | 0.99 |
| Body mass index (kg/m<sup>2</sup>) |  | 36.1 ± 5.1 | 31.5 ± 4.2 | **0.03** |
| HbA1c (%) |  | 7.1 ± 0.9 | 7.5 ± 0.9 | 0.26 |
| *Longitudinal lipid parameters* |  |  |  |  |
| Total cholesterol (mg/dL) | Baseline | 148.9 ± 11.0 | 164.7 ± 46.5 | 0.27 |
|  | 4 months | 140.0 [138.3–157.3] | 149.5 [140.0–169.3] | 0.43 |
|  | Delta | -7.0 [-12.0–8.5] | -5.0 [-15.3–20.0] | 0.92 |
| Triglycerides (mg/dL) | Baseline | 258.5 ± 55.3 | 285.2 ± 77.1 | 0.34 |
|  | 4 months | 253.4 ± 75.0 | 203.5 ± 72.3 | 0.12 |
|  | Delta | -5.1 ± 87.2 | -81.7 ± 69.8 | **0.03** |
| HDL-C (mg/dL) | Baseline | 34.5 ± 5.1 | 33.8 ± 4.5 | 0.80 |
|  | 4 months | 32.8 ± 5.9 | 33.6 ± 11.3 | 0.84 |
|  | Delta | -1.4 ± 4.6 | -0.2 ± 8.9 | 0.67 |
| LDL-C (mg/dL) | Baseline | 71.3 ± 12.4 | 82.2 ± 37.3 | 0.35 |
|  | 4 months | 69.0 [64.0–93.5] | 87.0 [87.0–99.8] | 0.35 |
|  | Delta | 5.5 ± 18.4 | 5.4 ± 29.6 | 0.99 |
| ApoB (mg/dL) | Baseline | 80.3 ± 14.8 | 89.7 ± 25.1 | 0.28 |
|  | 4 months | 81.0 ± 18.4 | 92.3 ± 18.3 | 0.14 |
|  | Delta | -1.8 [-7.9–5.8] | 6.75 [1.6–14.2] | 0.16 |
| Calculated VLDL (mg/dL) | Baseline | 47.0 [43.5–59.5] | 49.0 [45.0–63.0] | 0.64 |
|  | 4 months | 50.7 ± 15.0 | 40.8 ± 14.4 | 0.11 |
|  | Delta | -1.0 ± 17.4 | -14.4 ± 13.1 | **0.05** |
| ApoC3 (mg/dL) | Baseline | 15.2 ± 3.0 | 16.3 ± 2.6 | 0.36 |
|  | 4 months | 15.8 ± 4.6 | 12.2 ± 4.3 | 0.07 |
|  | Delta | 0.6 ± 4.7 | -4.0 ± 3.1 | **0.01** |

The serum proteome shift of pemafibrate-treated patients with HLD/DM reflected an attenuation of systemic inflammation. Principal component analysis (N=684 protein IDs) demonstrated a robust shift in the plasma proteome of patients treated with pemafibrate for four months as compared to their baseline (**Figure 6B**). In contrast, patients in the placebo group had very few differentially enriched proteins from their baseline to their 4-month follow-up: 20 in the placebo group vs. 207 in the pemafibrate group (**Figure 6C, Supplementary Figure S9A-C, Supplementary Table 17**). Differential enrichment analysis and pathway enrichment analysis in pemafibrate-treated patients demonstrated a consistent decrease in proteins related to immune cell response: monocyte activation, inflammation, and complement activation (C1QB/C, C1R, CHI3L1, PGLYRP2, PCYOX1, SAA2, HP), as well as an expected reduction in lipoprotein remodeling (APOC3, APOB, **Figure 6D-E, Supplementary Table S18**). Notably, the relative abundance of serum lipoprotein(a) (Lp(a)), a strong cardiovascular disease risk factor and pro-inflammatory chemotactic lipid, was reduced. However, because mice do not express Lp(a), its reduction is unlikely to be the primary driver of AS attenuation in our *in vivo* mouse model. Pemafibrate-treated patients had an increase in proteins related to ECM organization, exocytosis, cellular metabolism, and a reduction in coagulation. Of note, many of these proteins have been previously found to be anti-inflammatory and/or suppressive of macrophage adhesion and chemotaxis (ANGPTL4, DUSP3, TGFB1, GPX3, CHGA, GSN, VTN, APOA4), several through TLR activation and NFκB response^40,41^. MGP, a potent calcification inhibitor^42^, and proteins associated with lipoprotein clearance and HDL stability (APOA4, APOA2) were increased (**Figure 6D-E**). APOA4 reduces macrophage activation^43^. Several differentially enriched serum proteins that participate in macrophage activation and pro-inflammatory responses (S100A7, MMPs, CCL2) were increased in the serum. However, we found that in macrophages *in vitro*, these proteins were suppressed by serum from pemafibrate-treated HLD/DM Pt, suggesting a divergence between local and systemic inflammatory signatures. PCSK9 levels were also increased, which is a previously reported phenomenon in response to fibrates^44–46^. *GANAB*, a glucosidase regulating protein folding, was decreased by pemafibrate in both THP-1 macrophage-like cells and in the plasma proteome (**Supplementary Figure S9D)**.

### Pemafibrate-responsive proteomic features in human diseased valve tissue

Lastly, we sought to determine which pemafibrate-responsive proteomic features of our macrophage-VIC crosstalk *in vitro* studies were present in human aortic valve disease progression. Diseased human AV leaflets, which are comprised predominantly of VICs with local accumulation of pro-inflammatory macrophages in diseased regions, were obtained from patients undergoing surgical AV replacement, as we previously reported^47^ (N=18, **Figure 6F**). The valves were macroscopically dissected into non-diseased (ND), fibrotic (F), and calcified (C) regions. Region-annotated tissue proteomic analysis demonstrated a clear clustering of the calcified region proteome away from that of the fibrotic and non-diseased regions (**Figure 6G**). Using differential enrichment analysis, we identified proteins that were modified with calcific disease (n=1205, **Supplementary Table S19**) and performed overlap analysis to identify which of these features were pemafibrate-responsive in our *in vitro* experiments. Overlap signifies protein that were either i) increased in diseased regions in human AV tissue (C>ND, F>ND, C>ND; q<0.05) and then decreased *in vitro* by pemafibrate directly in THP-1 cells or indirectly by macrophage THP-1 CM in VICs, or ii) decreased in diseased regions in human AV tissue (C<ND, F<ND, C<ND, q<0.05) and increased by pemafibrate *in vitro* (VIC response to THP-1 cell CM, n=55; THP-1 cells, n=62; **Figure 6H-I** and **Supplementary Table S20)**. Directionality- and cell type-specific protein-protein interaction network clustering analysis identified that nonsense-mediated decay (GSPT1) and various protein translation regulation-related proteins (PDIA4, MYDGF) are decreased in calcified tissue and increased in response to pemafibrate in VICs (**Figure 6J, Supplementary Table S21**). Similarly, pemafibrate also rescued protein folding and translation-related proteins (RACK1) in THP-1 cells that were increased in calcified AS tissue. Proteins associated with derangements in cellular metabolism (NAD+ metabolism, carbohydrate metabolism) in AS were also increased by pemafibrate in the *in vitro* crosstalk data, suggesting these may be mechanistically translatable pathways of pemafibrate response therapeutically in humans. The *in vitro* derived pemafibrate responsive tissues also associated with calcification in human AV tissue mapped to cellular localization in the endoplasmic reticulum and specifically the ribosome, in alignment with perturbed protein translation and processing signatures (**Figure 6K**).

## Discussion

CAVD leads to AS, resulting in severe cardiovascular dysfunction, heart failure, and death. Pharmacological therapies for severe AS are lacking, and treatment is limited to invasive valve replacement. We evaluated the selective PPARα agonist pemafibrate as a therapeutic strategy for CAVD and AS. In a mouse AVWI model, pemafibrate attenuated AS and reduced AV calcification*. In vitro*, pemafibrate suppressed VIC calcification by modulating macrophage-derived secreted factors. Pemafibrate altered macrophage phenotype through anti-inflammatory and anti-chemotactic effects, both directly and via serum from pemafibrate-treated patients, thereby limiting inflammation-driven valvular calcification. These findings were supported by *post hoc* analysis of the PROMINENT randomized controlled trial, in which serum proteomics demonstrated suppression of inflammatory and innate immune pathways consistent with the in vitro results. Integration of human AS valve proteomics with the *in vitro* pemafibrate-responsive proteome identified disease pathways potentially rescued by pemafibrate in humans. Notably, this is the first study to demonstrate that a fibrate suppresses valvular calcification progression in an experimental AS model.

Both longitudinal echocardiography and valve leaflet imaging of calcification demonstrated that pemafibrate treatment attenuated features of AS and CAVD *in vivo.* Ldlr^-/-^ mice that underwent an AVWI model of AS treated with pemafibrate had a larger AV open diameter, lower peak transvalvular flow velocity, and higher AV area compared to untreated mice. EF also increased in the pemafibrate-treated HFD group at 15 weeks, suggesting these beneficial effects of pemafibrate on cardiac function may be partially due to reduced AV stenosis-driven pressure overload and/or favorable cardiac energetic changes. These findings are consistent with a recent fenofibrate clinical trial^48^. Mice fed a HFD exhibited a more severe CAVD phenotype as assessed by calcification, ECM remodeling enzymatic activity, and inflammation compared to ND controls, all of which were attenuated with pemafibrate. In the present study, pemafibrate reduced murine TG levels but did not alter total cholesterol levels, consistent with our PROMINENT trial sub-analysis. In humans, TG and calculated remnant cholesterol levels have been associated with AS risk in observational and Mendelian randomization studies^49,50^, and a higher TG/HDL-C ratio has been associated with the occurrence of CAVD^51^.

While TG-lowering effects of pemafibrate may constitute one of the mechanisms by which PPARα activation attenuated AV calcification in our *in vivo* studies, circulating TG levels showed no significant correlations with echocardiography parameters or the changes in valve inflammation, suggesting pleiotropic mechanisms of pemafibrate.

Accumulating evidence has established inflammation as a central driver of valvular calcification^25,52,53^. Macrophages are a characteristic feature of CAVD and a contributing cell type in the inception and progression of calcification^54,55^. Our *in vivo* and *in vitro* studies demonstrated anti-inflammatory effects of pemafibrate, consistent with our previous work showing that this compound shifts primary macrophages toward a less inflammatory phenotype, as assessed by single-cell profiling^20^. In the present study, *in vivo* pemafibrate administration in mice reduced macrophage accumulation (CD68^+^ cells), MMP activity, and calcification burden. *In vitro* data in the present study demonstrated the decreased expression of pro-inflammatory molecules in THP-1 macrophage-like cells treated with pemafibrate, either directly or indirectly via serum from pemafibrate-treated patients.

The secretion of pro-inflammatory cytokines, such as IFNγ, TNF-α, and IL-6, as well as pro-calcific extracellular vesicles by macrophages, have been implicated in promoting VIC-mediated calcification^14,15^. We demonstrated that exposing VICs to the secretome of macrophages was sufficient to induce calcification, and further that pre-treating the macrophages with pemafibrate attenuated this inflammation-driven calcification. VIC proteomic analysis of this macrophage-VIC crosstalk confirmed osteogenic and inflammatory activation and identified pemafibrate-responsive pathways related to ECM remodeling, protein translation, and bioenergetics as downstream secretome-mediated mechanisms. Integration of these data with proteomic profiles from human diseased aortic valve tissue confirmed that these pemafibrate-responsive pathways and candidate proteins identified *in vitro* are also dysregulated during human disease progression.

CAVD is an active inflammatory process involving immune cell accumulation, with both innate and adaptive immune responses contributing to disease progression^53^. We demonstrated that pemafibrate suppressed the gene expression of several chemokines in THP-1 macrophage-like cells. To capture the systemic response to pemafibrate, we exposed THP-1 monocytes and macrophage-like cells to serum from patients with HLD/DM treated with placebo or pemafibrate. Serum from pemafibrate-treated patients mitigated pro-inflammatory and pro-migratory effects on macrophages compared with placebo. Consistent with these findings, PPARα activation via fenofibrate has been shown to reduce monocyte/macrophage migration and suppress vascular and systemic inflammation in experimental atherosclerosis^56,57^. Together, our results indicate that pemafibrate exerts dual effects by attenuating macrophage inflammatory activation and potentially reducing their capacity to infiltrate the aortic valve during AS progression. In a *post hoc* sub-analysis from the PROMINENT clinical trial, serum proteomics revealed broad suppression of inflammatory pathways and proteins associated with monocyte activation following pemafibrate treatment, consistent with the *in vitro* findings.

The strengths of this study include the use of one *in vivo and* two *in vitro* models to demonstrate the attenuating effect of pemafibrate on AV inflammation and calcification. These findings are supported by the integration of serum and proteomics data from a clinical trial, as well as human diseased AV tissue. To better recapitulate human AS, we employed a modified AVWI model incorporating milder surgical valve injury on an AS-promoting Ldlr^⁻/⁻^ background with HFD feeding^25,54^. Pemafibrate exhibits greater PPARα potency than other fibrates, such as fenofibrate, as evidenced by its lower EC₅₀ value and higher subtype selectivity^19^. This selectivity for PPARα may be a contributing factor to the therapeutic outcomes observed in the AVWI model. The pathogenesis of CAVD involves several distinct phases, encompassing lipid accumulation, pro-inflammatory processes, fibrosis, and the differentiation of VICs to osteoblastic-like cells^58,59^. While the present study identifies a macrophage-VIC crosstalk axis as a mechanism underlying the anti-calcific effects of pemafibrate, it does not fully delineate the molecular pathways linking pemafibrate to the acceleration of valvular calcification. Our data support mechanisms of direct and indirect reduction of macrophage chemotaxis and release of pro-inflammatory molecules, such as cytokines or extracellular vesicles, as well as potentially systemic TG level reduction. However, the relative contribution of these factors, individually or in combination, to calcification progression remains to be determined.

In conclusion, these findings demonstrate for the first time the potential of fibrates, specifically pemafibrate, as a therapeutic strategy for AS. We identify a novel mechanism whereby pemafibrate mitigates experimental AS and AV calcification by reducing pro-inflammatory and pro-calcific activation of macrophages and modulating VIC-macrophage crosstalk. Furthermore, proteomic profiling using *post hoc* clinical trial serum, together with integration of human AS tissue data, substantiates these findings. Collectively, these results position pemafibrate as a potentially effective therapeutic agent for the treatment of AS that warrant future clinical investigation in patients with urgent unmet needs.

## Full Materials and Methods

### Aortic valve wire injury (AVWI) mouse model of AS

All mouse experiments were performed in compliance with the Institutional Animal Care and Use Committee at Beth Israel Deaconess Medical Center under animal protocol #014-2020 (Boston, MA, USA). Mice were maintained on a 12-hour light/dark cycle with food and water *ad libitum*. Male low-density lipoprotein receptor-deficient (Ldlr^-/-^) mice were purchased from the Jackson Laboratory (Bar Harbor, #002207). The animals were subjected to quarantine and subsequently maintained until 10-11 weeks of age. Ultrasound sonography was performed to acquire the basal blood velocities and AV open diameter in Ldlr^−/−^ mice, and the mice subsequently underwent AV wire injury (AVWI) surgery. After the recovery from the AVWI surgery (typically 3-5 days), the mice were randomly allocated to the pemafibrate treatment or non-treatment group. Mice were fed either normal diet (ND; Formulab Diet, LabDiet, #5008i) or a high-fat diet (HFD; Clinton/Cybulsky high-fat rodent diet with regular casein and 1.25% added cholesterol, Research Diets, Inc., #D12108COi) and each diet group was conducted in independent experiments. Each group had N=9-13 animals. Echocardiography was performed to evaluate cardiac function and aortic valve (AV) thickness at 8 and 15 weeks after AVWI. Multiphoton and confocal imaging were utilized to quantify AV calcification. Food intake and body weight were measured once every other week. No sham control arm was used in the study to minimize animal use as we have previously demonstrated the long-term AV profile of sham controls^7^ and described safety of pemafibrate at the dose provided in this study in Ldlr^-/-^ mice on an HFD^20^.

The surgical procedure for the AV stenosis mouse model was adapted from the original protocol described by Honda et al^25^ and as previously reported by our group^54^. In brief, Ldlr^-/-^ mice were anesthetized by inhalation of 1% isoflurane. The right carotid artery was exposed by dissection and ligated. A metal guide wire (ASAHI INTECC, MIRACLEBros 6 #AG14M060, diameter 0.36 mm) was inserted from the carotid into the left ventricle under echocardiographic guidance. The tip of the wire was positioned just below the AV level and moved forward and back 20 times and rotated 30 times to induce leaflet injury. The wire was removed, and the carotid artery was ligated. The peri-operative mortality rate of the procedure was below 10%. Animals that underwent the AVWI procedure were randomly assigned into four groups: normal diet (ND) control, ND containing 1.8 ppm pemafibrate, HFD control, and HFD containing 1.8 ppm pemafibrate. Diet containing 1.8 ppm pemafibrate is approximately equivalent to 0.2 mg/kg/day. According to pharmaceuticals and medical devices agency japan (PMDA) submission documents^60^, AUC of pemafibrate 0.4 mg/kg in humans was 23.305 ng*h/mL, and AUC of pemafibrate 0.2 mg/kg in mice was 26.4 ng*h/mL. We previously reported this dose of pemafibrate (0.2 mg/kg/day) mitigated vein graft lesion development and increased the patency of arteriovenous fistula failure^20^. All animals underwent AVWI and were fed ND or HFD, with or without pemafibrate for 15 weeks. Cardiac function was monitored using echocardiography prior to AVWI and after AVWI at 8 and 15 weeks. AV open diameter and flow peak velocity were measured with a pulse wave Doppler and echocardiography (MS250 transducer, VEVO3100, Fujifilm VisualSonics). Briefly, mice were anesthetized using 1.5% isoflurane with an oxygen flow rate of 1.0 L/min on a thermal platform set to 40°C. The right parasternal long-axis view in B-mode was utilized for measures of AV open diameter and flow peak velocity. The left parasternal long axis view in pulse-wave and color Doppler was used to measure transvalvular blood velocities. Doppler measures of blood velocity were done with the intercept angle less than 60° between the vessel and ultrasound beam. Each parameter was measured as the average in three cardiac cycles. Aortic valve area (AVA) was measured by continuity equation.

### Visualization of calcification and MMP activity in AV

At 15 weeks after surgery, two near-infrared fluorescent probes, OsteoSense680 and MMPSense750 (PerkinElmer Inc. #NEV10020EX and #NEV10168, respectively), were injected intravenously 24-hours before sacrifice. Two imaging probes (2 nmol/100 μL, each probe) were injected into the mouse via tail vein. OsteoSense is a bisphosphonate conjugated to a fluorophore, which binds specifically to calcium-phosphate crystal in calcification. MMPsense is a fluorophore conjugated to a peptide substrate for matrix metalloproteases (MMPs) that fluoresces when cleaved allowing for local detection of MMP activity. At sacrifice, blood was collected from vena cava, then animals were perfused with saline before harvesting organs. AV leaflets were micro-dissected and subjected to imaging analysis to visualize calcification (Osteosense: filter setting; 675 excitation/720 emission) and MMP activity (MMPsense: 745 excitation/800 emission) presence in the leaflet. Images were captured with a fluorescent microscope (Olympus MVX10) using the filter with CY5 for OsteoSense680 or CY7 for MMPsense750 (exposure around 1 to 2 sec, gain x1) and CellSens imaging software (Olympus Corporation. Tokyo, Japan). Signal intensity was quantified using Fiji ImageJ software (version: 2.14.0/1.54f).

### Mouse plasma biomarker measurement

Blood was collected from vena cava using a heparinized syringe with 5,000 units/mL heparin (Patterson Veterinary Supply, Inc., #07-892-8971) into a 1.5 mL tube, followed by centrifugation (4°C, 1000xg, 15 min) to collect plasma. Total cholesterol or triglyceride levels in plasma were measured using commercial kits (Sigma Aldrich, #MAK043 and #MAK266, respectively) according to the manufacturer’s protocol. Plasma calcium and phosphate concentration was measured using QuantiChrom Calcium Assay Kit or QuantiChrom Phosphate Assay Kit (BioAssay systems, #DICA-500 and #DIPI-500, respectively) according to the manufacturer’s protocol.

### Histopathology

AVs were embedded in OCT compound (Fisher Healthcare, #23-730-571). Cryosections (7 μm) were stained with hematoxylin and eosin (H&E). AV thickness was measured with H&E sections. Thickness at four longitudinal points of each leaflet was measured using ImageJ software and the average value was used to calculate mean valve thickness of each animal. Macrophage infiltration of AVs was assessed using CD68 antibody immunohistochemistry (IHC) (Abcam, #ab5344, 1:100). IL-12 and S100A9 IHC was performed using anti-IL-12 (Novus Biologicals, #NB6001443, 1:100) and anti-S100A9 (Proteintech, #26992-1-AP, 1:100) antibodies, respectively. The primary antibody was incubated at 4°C overnight, followed by incubation with the secondary antibody (Alexa Fluor 594 anti-Rat, Abcam, #a21209, 1:200 dilution for CD68 and IL-12; Alexa Fluor 488 anti-Rabbit, Abcam, #a21206, 1:200 dilution for S100A9) at room temperature for 45 minutes in a humidity chamber. Nuclear counterstain was performed using NucBlue (Invitrogen, #R37606), followed by using VectaShield anti-fade mounting media (Vector Laboratories, #H-1700) for 1-2 hours at room temperature. H&E staining and immunohistochemistry images were acquired using NIS elements software (Nikon, version 5.10.01. 64-bit) and analyzed using Fiji ImageJ software (version 2.14.0/1.54f). In the histopathological analysis of calcification (OsteoSense680 imaging) and IL-12 and S100A9 IHC, a consistent threshold was applied to all images for each staining protocol, utilizing negative control staining as a reference. The signal area across the entire valve section was then quantitatively assessed using Fiji ImageJ software. The data was normalized by the DAPI positive area. CD68^+^ cells in the valve leaflet area were counted and normalized by the length of the leaflet.

### Pemafibrate-treated human serum samples

Human serum samples were obtained from PROMINENT clinical trial, a randomized controlled trial whereby adult participants with type 2 diabetes and hypertriglyceridemia received 0.2 mg of pemafibrate or placebo twice daily for four months^24^. Serum was obtained from 24 patients (12 placebo, 12 pemafibrate treatment) collected two weeks after the initiation of the clinical trial (Visit 1). For the chemotaxis assay, serum from patients administered placebo or pemafibrate was pooled with equal volume from each donor. Pooled serum was added into Roswell Park Memorial Institute-1640 medium (RPMI-1640; Corning, #MT10040CV) at 5% volume concentration.

### Cell culture

Human primary valvular interstitial cells (VICs) were isolated from human AV leaflets from patients undergoing surgical AV replacement as part of their clinical care. The study protocol was approved by the Institutional Review Board and Human Research Committee at Brigham and Women’s Hospital (#2011P001703). VICs were isolated from AV leaflets using collagenase digestion as previously reported^61^. In brief, both sides of the leaflet were scratched by razor blade to remove endothelial cells. Leaflets were cut into 1-2mm^3^ pieces and digested with 1mg/mL collagenase type I (Sigma Aldrich, #C5894) in high-glucose Dulbecco’s Modified Eagle Medium (DMEM; ThermoFisher, #11965118) for 1 hour at 37°C. Pieces were rinsed with DMEM and then further digested with 1 mg/mL collagenase type-I for 3 hours. VICs were isolated from the second digest solution via centrifugation (500xg, 4°C) and plated in 75 cm^2^ culture flasks. Isolated VICs were cultured in normal media (NM: high-glucose DMEM supplemented with 10% heat-inactivated fetal bovine serum (FBS; Avantor, #97068-085) and 1% penicillin/streptomycin (P/S; ThermoFisher, #15140163), 1mM sodium pyruvate) until cells were >90% confluent. Cells were used for experiments at passages 3-5 and were plated for each experiment at 5.26 x10^4^ cells/cm^2^. For experiments, the VICs were cultured in NM, osteogenic media (OM; DMEM with 10% FBS, 1% P/S, 10 nM dexamethasone, 10 mM β-glycerol phosphate, and 100 mM L-ascorbate phosphate), or NM containing 50% RPMI-based THP-1-derived conditioned media (CM) for at least 21 days to assess calcification phenotype. Media was replaced every 3-4 days.

THP-1 monocytes were purchased from ATCC (#TIB-202) and maintained in complete RPMI-1640 medium containing 10% FBS and 1% P/S. THP-1 monocytes were differentiated into THP-1 macrophage-like cells using phorbol 12-myristate 13-acetate (PMA, Sigma, #P1585; 100 ng/mL). THP-1 macrophage-like cells were polarized to a pro-inflammatory M1 state, M(LPS+IFNγ), through stimulation with lipopolysaccharide (LPS, InvivioGen, #tlrl-smlps; 10 pg/mL) and interferon-γ (IFNγ, R&D Systems, #285-IF-100; 20 ng/mL) for 48 hours. Either non-polarized THP-1 macrophage-like cells, M(-), or M(LPS+IFNγ) were cultured with pemafibrate (10 μM), control human serum (Gemini Bio-Products, #100-512), or serum from PROMINENT participants (final concentration: 5%) for 48 hours, followed by gene expression analysis.

### THP-1 conditioned media (CM) preparation

THP-1 monocytes were subcultured in T75 flasks at 6.7 x 10^3^ cells/cm^2^. To induce differentiation into THP-1 macrophage-like cells, THP-1 monocytes were plated on 150 mm dishes (Falcon, #353025, 4 x 10^5^ cells/cm^2^) with phorbol 12-myristate 13-acetate (PMA, 100 ng/mL) and incubated for 48 hours, followed by a resting culture in RPMI-1640 media without PMA for 24 hours. Subsequently, cells were cultured with or without 0.1, 1, or 10 μM pemafibrate, according to the previous report^62^, for 48 hours, and the conditioned media (CM) was collected at 24-hour intervals by replacing Pema-contained complete media. CM was centrifuged at 300xg for 5 minutes to remove cellular debris. NM containing 50% CM exposed to VICs for calcification assay. Media without exposure to cells was used as control media (NM). Aliquots were stored at -30 °C.

### Chemotaxis assay

The chemotaxis assay was performed according to the manufacturer’s protocol (Sartorius). Briefly, THP-1 monocytes were differentiated to THP-1 macrophage-like cells with 100 ng/mL PMA for 24 hours, followed by 24 hours of resting culture without PMA. THP-1 macrophage-like cells were detached using Accutase (STEMCELL Technologies, #07920) and subsequently plated 3,000 cells per well in the top chamber well of IncuCyte Clearview 96-well plate for chemotaxis (Sartorius, #4600). RPMI-1640 media with 1% P/S and 5% serum from placebo- or pemafibrate-treated PROMINENT participants were placed in the bottom chamber (200 μL/well). The chemotaxis plate was subsequently placed into the IncuCyte S3 (Sartorius) for imaging the live cell migration for 12 hours. The data were analyzed using the Live-Cell Imaging and Analysis System module.

### Gene expression analysis

RNA was extracted from either M(-) or M(LPS+IFNγ) 48 hours post-exposure to 10 μM Pema, 5% placebo serum-, 5% Pemafibrate serum-, or 5% control serum-containing complete RPMI-1640 media, utilizing TRI reagent (Sigma, #T9424). Subsequently, 1 μg of total RNA from each sample underwent reverse transcription using qScript cDNA Synthesis Kit (Quantabio, #95047). The synthesized cDNA was then subjected to pre-amplification (14 cycles) using PerfeCTa® PreAmp SuperMix (Quantabio, #95146) in accordance with the manufacturer’s protocol. Gene expression was quantified by Fluidigm BioMark HD Real-Time PCR and its analysis module (Standard Bio Tools) using the probes listed in Supplementary Table S1.

### Alizarin red staining

To assess the deposition of calcium, VICs were fixed with 4% paraformaldehyde for 10 minutes, then stained with 2% alizarin red stain (Lifeline Cell Technology, #CM-0058) for 15 minutes. Wells were washed three times with water and images were captured from the bottom side of the plate. To quantify alizarin red staining, 100uL of 100 mmol/L Cetylpyridinium chloride (Sigma, #C0732-100G) was added to each well and gently rotated at room temperature for 30 minutes. The absorbance of the resultant solution was read at 540nm for quantification.

### Cell lysate proteomics

Monolayer cultured cells were rinsed twice with PBS, and then cell lysates were harvested using RIPA buffer supplemented with protease and phosphatase inhibitors (Roche, cOmplete Mini tablets, #4693159001, and phosSTOP, #4906845001). Lysate samples were centrifuged at 16,000 g for 10 min at 4°C. The supernatant was collected and transferred to a new tube and stored at -80°C until further analysis. The protein concentration was determined using Pierce BCA protein assay (ThermoFisher, #PI23227). 5.3-7.5 ug (VIC) or 23ug (THP-1) protein was used for proteomics preparation. Peptides for MS injection from cell lysates were prepared using the PreOmics iST kit protocol using the PreON sample preparation robot. In LYSE buffer, samples were heated at 95 °C for 10 minutes. Protein samples were digested using PreOmics provided Lys-C/trypsin DIGEST solution for 2 hours at 37 °C. Samples were speed-vacuumed at 45°C only until the peptides were dried and were subsequently stored at -80°C until LC-MS/MS sequencing. 42 uL of LC-LOAD solution (PreOmics) was added to the dried peptides. Samples were centrifuged at 16,000 G for 1 minute and then 40 uL were aspirated into a new tube and 4 μL of a 5-fold dilution was injected into the mass spectrometer.

Peptide samples were injected into a quadrupole Orbitrap Exploris 480™ coupled to a Vanquish™ Neo UHPLC system (Thermo Fisher Scientific). The peptides were passed through a dual column setup with a PepMap™ Neo C18 Trap Cartridge, 5 μm, 300 μm X 5 mm (Thermo Fisher Scientific, # 174500), and an Easy-Spray™ PepMap™ Neo C18 Column, 2 μm, 75 μm X 150 mm (Thermo Fisher Scientific, # ES75150PN). The column was heated at a constant temperature of 45 °C. The gradient flow rate was 300 nL/min from 5 to 21% solvent B (95% acetonitrile /0.1% formic acid in mass spectrometry-grade water) for 50 min, 21 to 30% solvent B for 10 min, and another 15 min of a 95%-5% solvent B sawtooth wash. Solvent A was 0.1% formic acid in the water. The mass spectrometry analysis was performed using data-dependent acquisition (DDA) and operated in positive mode. Spray voltage was set to 2,000 V, funnel RF level at 40, and heated capillary temperature at 275 °C. MS1 resolution was set to 120K, and scan range was m/z 375–1500. The ions with charge state of 2-6 were selected for MS2 fragmentation, dynamic exclusion duration was 60 seconds, and intensity threshold was set to 50,000. The total cycle time of the data dependent mode was set to 3 s. MS2 resolution set to 60,000, isolation window was 1.2 m/z, AGC target was standard, maximum injection time was auto, and normalized HCD collision energy was steps of 24, 26 and 28%.

The acquired peptide spectra were searched with Proteome Discoverer package (PD, Version 2.5) using the SEQUEST-HT search algorithm against the Human UniProt database (42,421 entries, downloaded May 2025). Cell types were searched separately. Trypsin (full) was set as the digestion enzyme, allowing up to 2 missed cleavages and a minimum peptide length of 6 amino acids. Oxidation (+15.995 Da) of methionine; and acetylation (+42.011 Da) of the N-terminus, were set as variable modifications. Carbamidomethylation (+57.021 Da) of cysteine was set as a static modification. Spectral search tolerances were 10 ppm for the precursor mass and 0.02 Da for HCD spectra. Peptides were filtered based on a false discovery rate of 1.0%, which was calculated using Percolator (target/decoy method, separate databases), provided by Proteome Discover and peptides were filtered based on a 1.0% FDR. Quantification utilized unique peptides (those assigned to a given Master protein group and not present in any other protein group) and razor peptides (peptides shared among multiple protein groups). Razor peptides were used to quantify only the protein with the most identified peptides and not for the other proteins they are contained in. The “Feature Mapper” was enabled in PD to identify peptide precursors that may not have been sequenced in all samples but were detected in the MS1. The chromatographic spectra were aligned while allowing for a maximum retention time shift of 10 minutes, mass tolerance of 10 ppm, and a minimum signal-to-noise ratio of 5. Chromatographic intensities were used to establish precursor peptide abundance. Peptide abundance was normalized by total peptide amount. Each protein intensity was calculated using the sum of its peptide intensities. A minimum of at least 2 unique peptides for each protein was required for the protein to be included in the analyses.

### Conditioned media proteome

The proteome of the THP-1-derived CM was measured. For application to the VICs, all media was pooled and then stored at -20C in aliquots, thus there is a single replicate per conditioned media condition. The CM was diluted two-fold in water and ultrafiltered to 50% volume using Pierce Concentrator PES 3K MWCO 0.5mL (Thermo, #88512). Peptides were prepared using PreOmics iST kit protocol with an input of 25uL of media and 25uL of 2X LYSE buffer. Protein samples were digested using PreOmics provided Lys-C/trypsin DIGEST solution for 2 hours at 37 °C. Subsequent peptides were quantified with the ThermoScientific NanoDrop and diluted to 75 ng/uL with LC-LOAD buffer. Compared to the cell lysate samples, minor modifications were made. Specifically, instead of SEQUEST-HT search algorithm node the Fixed PSM node was used to allow for more potential contaminants to be identified.

### Serum proteomics

Serum samples collected from patients enrolled in the PROMINENT clinical trial at the 4-month time-point were prepared for serum proteomics including patients from both the placebo (n=12) and pemafibrate group (n=12). Peptides were prepared as per protocol using the PreOmics ENRICH-iST kit designed for serum and plasma. As per protocol, 50uL of serum was partially depleted of highly abundant serum proteins using magnetic beads prior to peptide preparation to allow for deeper proteome sequencing.

Tryptic peptides were analyzed using the quadrupole Orbitrap Exploris 480™ coupled to a Vanquish™ Neo UHPLC system (Thermo Fisher Scientific). The peptides (400 ng on column) were fractionated using a dual column set-up: a PepMap™ Neo C18 Trap Cartridge, 5 μm, 300 μm X 5 mm (Thermo Fisher Scientific, Cat# 174500); and an Easy-Spray™ PepMap™ Neo C18 Column, 2 μm, 75 μm X 150 mm (Thermo Fisher Scientific, Cat# ES75150PN). The column was heated at a constant temperature of 45 °C. The gradient flow rate was 300 nL/min from 5 to 21% solvent B (95% acetonitrile /0.1% formic acid in mass spectrometry-grade water) for 60 min, 21 to 30% solvent B for 10 min, and another 15 min of a 95%-5% solvent B sawtooth wash. Solvent A was 0.1% formic acid in water. The mass spectrometry analysis was performed using data-independent acquisition (DIA). MS1 scans were acquired in profile mode and MS2 scans in centroid mode. Spray voltage was set to 2,000 V, funnel RF level at 50, and heated capillary temperature at 275 °C. MS1 were with 120K resolution and mass scan range was set to m/z 400–900. The AGC target was set to standard with a maximum injection time of 25 ms. MS2 spectra were acquired with a precursor isolation range of m/z 400−900 divided into 4 Th windows with an overlap of 1 Da. MS2 resolution was set to 60 K and AGC target value for fragment spectra was set at 1000% with a maximum injection time of auto. HCD collision energy was set to 26%.

The acquired peptide spectra were searched with the Proteome Discoverer package (PD, Version 3.2) using the CHIMERYS node with the INFERYS 4.7.0 prediction model. The spectra were queried against the Uniprot human database (downloaded May 2025; 42,421 entries). Trypsin/P was set as the digestion enzyme, allowing up to 2 missed cleavages and a peptide length of 7-30 amino acids (1-6 peptide charge). Oxidation (+15.995 Da) of methionine was set as a variable modification, and carbamidomethylation (+57.021 Da) of cysteine was set as a static modification. The fragment mass tolerance was 20ppm. The FDR for PSM confidence was 0.01 and 0.05, for strict and relaxed confidence, respectively. The quantification type was set to the default setting, MS2 Apex (Quan in all files). Quantification utilized unique peptides and razor peptides.

Razor peptides were used to quantify only the protein with the most identified peptides and not for the other proteins they are contained in. Peptide abundance was normalized by total peptide amount, and each protein intensity was calculated using the sum of its peptide intensities. Only protein groups with high FDR confidence were used in analysis.

### Valve leaflet tissue proteomics

Publicly available RAW mass spectrometry files from PRIDE Archive (PXD035538) with matched non-diseased, fibrotic, and calcified regions (N=54) were searched and annotated as per the “Cell Lysate Proteomics” section above. Valve leaflet samples were obtained, dissected, prepared for proteomics, and injected as described previously^47^. In brief, 18 diseased tricuspid AV leaflet samples were obtained from consented patients with aortic stenosis undergoing surgical valve replacement at Brigham and Women’s Hospital under approved protocol 2011P001703 and were grossly dissected into non-diseased, fibrotic, and calcified regions. 77.8% of patients were male and aged 69±7 and other demographics are available in the original publications^47,63^.

### Proteome analysis

The quantified proteins were exported from Proteome Discoverer and pre-processed using Perseus^64^ (V2.0.11 for VIC and human tissue, and V2.1.5.0 for THP-1 cells and serum). Proteins were removed from analysis if they were not present in 70% of samples in at least one experimental group, the recommended pre-processing cut-off^64^. The datasets were median (VIC, THP-1, serum) or Z-score (tissue) normalized, log2 transformed, and missing data was imputed using a Gaussian distribution within each sample with a 2 standard deviation downshift within a 0.2 standard deviation range. For VIC cell lysates (5 donors), serum proteomics (24 participants), and tissue proteomics (18 donors), variance from donors or participants was removed using a generalized linear model in Qlucore 3.10. Experiment-specific statistical testing and hierarchical clustering for each figure is described in each figure caption in detail and was performed using Perseus with a false discovery rate corrected p-value threshold (referred to in text as a q-value) of 0.05. Proportional Venn diagrams were generated using DeepVenn (arXiv:2210.04597, adapted from BioVenn^65^ by the author). Pathway enrichment analysis and protein-protein interaction networks were performed using StringDB V12.0^66^ (accessed May 2025-Sept 2025). Pathway-protein networks were generated using EnrichrKG^67^ and Reactome 2022. Networks were visualized using Cytoscape V3.10.2.

### Statistical Analysis

All data are presented as mean ± SEM. Two-way ANOVA followed by Bonferroni *post hoc* test or Tukey’s multiple comparison test, one-way ANOVA followed by Dunnett’s multiple comparison test, or Student *t*-tests were performed for statistical analysis by Prism 10 software (GraphPad software). *p<0.05, **p<0.01, ***p<0.001. For proteomics data analysis, specific statistical tests are outlined in figure captions.

## Acknowledgments

The authors thank the members and management team in the Center for Interdisciplinary Cardiovascular Sciences for their contributions. Schematic figures 1A, 3A, 4A, 5A, 5F, 5J, 6A, and 6H were created using BioRender.com.

## Author Contributions

YN, MT, TT, MA, and EA conceptualized the study. YN, MT, and EA authored the initial draft of the research manuscript. YN, MT, and TT were responsible for the analysis and interpretation of the collected data. YN, MT, TT, MB, AL, TK, SI, RG, KP, RI, TO, and YS participated in the acquisition of requisite information and data for the investigation. AP, PL, and PR shared PROMINENT clinical trial sample and information. KP conducted histological examinations. YN, MT, MB, SAS, MA, and EA composed the preliminary version of the paper. Each set of initials represents a distinct author or contributor to the research project.

## Supplementary Materials

**Supplementary Figure S1: Body weight and heart weight in AVWI mice with pemafibrate treatment at week 15**

**(A)** Body weight (g), **(B)** Heart weight (g), and **(C)** Relative values of heart weight/body weight in AVWI model at 15 weeks after surgery. Mean±SEM. N=10-12/group. Ordinary two-way ANOVA followed by Bonferroni post hoc test performed for statistical analysis between control and pemafibrate in ND and HFD. *p<0.05.

**Supplementary Figure S2: Plasma calcium decreased and phosphate level did not change with pemafibrate in AVWI mice at week 15**

**(A)** Total calcium (mmol/L) and **(B)** phosphate (μmol/L) in plasma in AVWI model at 15 weeks after surgery. Mean±SEM. N=10-12/group. Ordinary two-way ANOVA followed by Bonferroni *post hoc* test performed for statistical analysis between control and pemafibrate in ND and HFD. *p<0.05.

**Supplementary Figure S3: Pemafibrate suppressed aortic valve leaftlet calcification in HFD group**

Osteosense680 signal area in the aortic valve 15 weeks after AVWI (Blue: DAPI, White: OsteoSense680). Mean±SEM. N=10-12/group. Scale bar = 100μm. Ordinary two-way ANOVA followed by Bonferroni *post hoc* test performed for statistical analysis between control and pemafibrate in ND and HFD. *p<0.05.

**Supplementary Figure S4: Echocardiography parameters did not correlate with plasma triglyceride level**

Plasma triglyceride levels plotted against echocardiography parameters. The parameters were acquired at the 15-week time point in high-fat diet-fed group. Linear regression plotted with 95% confidence bands of the best-fit line.

**Supplementary Figure S5: Plasma triglyceride levels did not correlate with aortic valve inflammation markers or calcified aortic valve area using histopathology**

Plasma triglyceride levels plotted against inflammation markers (IL-12 and S100A9) or calcification (Osteosense) in the aortic valve leaflet. The parameters were acquired at the 15-week time point in high-fat diet-fed group. Linear regression plotted with 95% confidence bands of the best-fit line.

**Supplementary Figure S6: Pemafibrate did not directly inhibit calcification of human primary VICs**

**(A)** Schematic of *in vitro* assay in VICs treated with pemafibrate. **(B)** Representative images of alizarin red staining in VICs (NM, normal media; OM, osteogenic media). **(C)** Donor-specific quantification of alizarin red staining in human VICs. The plots represented the values from independent duplicate wells. Ordinary one-way ANOVA followed by Dunnett’s multiple comparison test.

**Supplementary Figure S7: Conditioned media from THP-1 macrophage-like cells did not change ALP activity in VICs, regardless of pemafibrate treatment**

ALP activity value was normalized by protein amount. N=6 donors per group.

**Supplementary Figure S8: 5% of differentially enriched proteins in the VIC proteomic analysis proteins are also identified in the secretome and reflect proteins that may in-part or wholly be derived from THP-1 cells**

**(A)** Overlap of protein IDs found in THP-1 conditioned media (secretome proteome, n=604) and the experimental NM control (n=265), containing only fetal bovine serum and media. The unique secretome proteome (n=386, Supplementary Table S10) **(B)** Overlap of protein IDs from the secretome compared to all differentially enriched proteins in the pemafibrate analysis presented in Figure 5 (n=223). 23 overlap with the entire secretome including FBS-derived and 11 (bolded in in Figure) are only identified in CM. Experimental VICs were treated with pooled conditioned media, thus n=1 for each condition (NM, CM, CM + Pema 0.1, 1, 10 μM)

**Supplementary Figure S9: Serum proteomic analysis of placebo group from PROMINENT**

**(A)** Principal component analysis serum proteome (684 IDs) of patients from the placebo arm at baseline and following 4 months. Effect of participant variability removed using a generalized linear model. **(B)** Differential enrichment analysis comparing baseline to 4 months following pemafibrate using a T-test. Highlighted in red and blue and red are proteins with FDR q<0.05. **(C)** Overlap of differentially enriched proteins in placebo and pemafibrate arm. **(D)** Z-score normalized protein abundance in placebo arm plotted against the pemafibrate arm of the differentially enriched proteins in the pemafibrate-arm (Figure 6E). Subsets of proteins selected for further assessment with large change in pemafibrate Z-score normalized abundance (>0.6) and small change in the placebo arm (<0.1).

**Supplementary Table S1: qPCR probe list**

**Supplementary Table S2: Echocardiography parameters in AVWI model**

**Supplementary Table S3: Gene ID list of differentially enriched proteins between NM and CM. ANOVA followed by *post hoc* Tukey HSD test with q<0.05.**

**Supplementary Table S4: All enriched Reactome pathways (q<0.05) from gene IDs listed in Table S3.**

**Supplementary Table S5: Gene ID list of differentially enriched proteins between CM and any pemafibrate group (0.1, 1, 10μM pemafibrate) separated by directionality at each dose. ANOVA followed by *post hoc* Tukey HSD test with q<0.05.**

**Supplemental Table S6: Gene ID list of any differentially enriched protein between CM and any pemafibrate group separated by directionality. ANOVA followed by *post hoc* Tukey HSD test with q<0.05.**

**Supplementary Table S7: All enriched Reactome pathways (q<0.05) from gene IDs listed in Table S6.**

**Supplementary Table S8: Dose-dependent proteomic clustering in response to conditioned media. Unabridged k-means clustering annotations from StringDB protein-protein interaction network.**

**Supplementary Table S9: Top 8 Gene Ontology Biological Processes pathway annotations from pathway enrichment analysis and network generation using EnrichrKG.**

**Supplementary Table S10: Gene ID list of unique secretome proteome excluding the FBS-NM control.**

**Supplementary Table S11: Reactome and Gene Ontology Cellular Component pathway enrichment analysis of gene IDs in Supplementary Table S10.**

**Supplementary Table S12: Reactome pathway enrichment analysis of gene IDs (n=23) overlapping in differential enrichment analysis and secretome.**

**Supplementary Table S13: Differential enrichment analysis of THP-1 cells treated with pemafibrate or vehicle. T-test.**

**Supplementary Table S14: Gene ID list of differentially enriched proteins (q<0.05) fromTHP-1 cells treated with pemafibrate or vehicle.**

**Supplementary Table S15: Unabridged Reactome pathway enrichment analysis of gene ID list in Supplementary Table S14.**

**Supplementary Table S16: Reactome functional enrichment analysis of comparison and fold change in Supplementary Table S13.**

**Supplementary Table S17: Gene ID list of proteins differentially enriched (q<0.05) between baseline and 4-month time points in the placebo and pemafibrate arm separately.**

**Supplementary Table S18: Unabridged list of top 5 enriched gene ontology Biological Processes generated by EnrichrKG.**

**Supplementary Table S19: Differentially enriched proteins in human aortic valve tissue proteome dataset. ANOVA followed by *post hoc* Tukey HSD test with q<0.05.**

**Supplementary Table S20: Gene ID list from overlap analysis of differentially enriched proteins in tissue with disease progression, and those identified to be pemafibrate-responsive in THP-1 and VICs**

**Supplementary Table S21: Unabridged network annotations for stringDB PPIs clustered using Markov cluster algorithm with a granularity parameter of 4.**

## References

1. Coffey S, Cox B, Williams MJ. The prevalence, incidence, progression, and risks of aortic valve sclerosis: a systematic review and meta-analysis. J Am Coll Cardiol. 2014;63:2852–2861. doi: 10.1016/j.jacc.2014.04.018

2. Osnabrugge RL, Mylotte D, Head SJ, Van Mieghem NM, Nkomo VT, LeReun CM, Bogers AJ, Piazza N, Kappetein AP. Aortic stenosis in the elderly: disease prevalence and number of candidates for transcatheter aortic valve replacement: a meta-analysis and modeling study. J Am Coll Cardiol. 2013;62:1002–1012. doi: 10.1016/j.jacc.2013.05.015

3. Mengi S, Januzzi JL, Jr., Cavalcante JL, Avvedimento M, Galhardo A, Bernier M, Rodes-Cabau J. Aortic Stenosis, Heart Failure, and Aortic Valve Replacement. JAMA Cardiol. 2024;9:1159-1168. doi: 10.1001/jamacardio.2024.3486

4. Kraler S, Blaser MC, Aikawa E, Camici GG, Luscher TF. Calcific aortic valve disease: from molecular and cellular mechanisms to medical therapy. Eur Heart J. 2022;43:683–697. doi: 10.1093/eurheartj/ehab757

5. Blaser MC, Back M, Luscher TF, Aikawa E. Calcific aortic stenosis: omics-based target discovery and therapy development. Eur Heart J. 2024. doi: 10.1093/eurheartj/ehae829

6. Abdelbaky A, Corsini E, Figueroa AL, Subramanian S, Fontanez S, Emami H, Hoffmann U, Narula J, Tawakol A. Early aortic valve inflammation precedes calcification: a longitudinal FDG-PET/CT study. Atherosclerosis. 2015;238:165–172. doi: 10.1016/j.atherosclerosis.2014.11.026

7. Iqbal F, Schlotter F, Becker-Greene D, Lupieri A, Goettsch C, Hutcheson JD, Rogers MA, Itoh S, Halu A, Lee LH, et al. Sortilin enhances fibrosis and calcification in aortic valve disease by inducing interstitial cell heterogeneity. Eur Heart J. 2023;44:885–898. doi: 10.1093/eurheartj/ehac818

8. Hutcheson JD, Goettsch C, Bertazzo S, Maldonado N, Ruiz JL, Goh W, Yabusaki K, Faits T, Bouten C, Franck G, et al. Genesis and growth of extracellular-vesicle-derived microcalcification in atherosclerotic plaques. Nat Mater. 2016;15:335–343. doi: 10.1038/nmat4519

9. Li G, Qiao W, Zhang W, Li F, Shi J, Dong N. The shift of macrophages toward M1 phenotype promotes aortic valvular calcification. J Thorac Cardiovasc Surg. 2017;153:1318–1327 e1311. doi: 10.1016/j.jtcvs.2017.01.052

10. Turner ME, Bartoli-Leonard F, Aikawa E. Small particles with large impact: Insights into the unresolved roles of innate immunity in extracellular vesicle-mediated cardiovascular calcification. Immunol Rev. 2022;312:20–37. doi: 10.1111/imr.13134

11. New SE, Goettsch C, Aikawa M, Marchini JF, Shibasaki M, Yabusaki K, Libby P, Shanahan CM, Croce K, Aikawa E. Macrophage-derived matrix vesicles: an alternative novel mechanism for microcalcification in atherosclerotic plaques. Circ Res. 2013;113:72–77. doi: 10.1161/CIRCRESAHA.113.301036

12. Rogers MA, Buffolo F, Schlotter F, Atkins SK, Lee LH, Halu A, Blaser MC, Tsolaki E, Higashi H, Luther K, et al. Annexin A1-dependent tethering promotes extracellular vesicle aggregation revealed with single-extracellular vesicle analysis. Sci Adv. 2020;6. doi: 10.1126/sciadv.abb1244

13. Li XF, Wang Y, Zheng DD, Xu HX, Wang T, Pan M, Shi JH, Zhu JH. M1 macrophages promote aortic valve calcification mediated by microRNA-214/TWIST1 pathway in valvular interstitial cells. Am J Transl Res. 2016;8:5773–5783.

14. Grim JC, Aguado BA, Vogt BJ, Batan D, Andrichik CL, Schroeder ME, Gonzalez-Rodriguez A, Yavitt FM, Weiss RM, Anseth KS. Secreted Factors From Proinflammatory Macrophages Promote an Osteoblast-Like Phenotype in Valvular Interstitial Cells. Arterioscler Thromb Vasc Biol. 2020;40:e296–e308. doi: 10.1161/ATVBAHA.120.315261

15. Song X, Song Y, Ma Q, Fang K, Chang X. M1-Type Macrophages Secrete TNF-alpha to Stimulate Vascular Calcification by Upregulating CA1 and CA2 Expression in VSMCs. J Inflamm Res. 2023;16:3019–3032. doi: 10.2147/JIR.S413358

16. Tagzirt M, Rosa M, Corseaux D, Vincent F, Vincentelli A, Daoudi M, Jashari R, Staels B, Van Belle E, Susen S, et al. Modulation of inflammatory M1-macrophages phenotype by valvular interstitial cells. J Thorac Cardiovasc Surg. 2023;166:e377–e389. doi: 10.1016/j.jtcvs.2022.08.027

17. Baddour T, Ninh VK, Gorashi RM, Pena R, King KR, Aguado BA. Spatial transcriptomic profiling of the human aortic valve reveals cellular sex differences near sites of calcification. bioRxiv. 2025. doi: 10.1101/2025.08.20.671175

18. Felix Velez NE, Tu K, Guo P, Reeves RR, Aguado BA. Secreted Cytokines From Inflammatory Macrophages Modulate Sex Differences in Valvular Interstitial Cells on Hydrogel Biomaterials. J Biomed Mater Res A. 2025;113:e37885. doi: 10.1002/jbm.a.37885

19. Raza-Iqbal S, Tanaka T, Anai M, Inagaki T, Matsumura Y, Ikeda K, Taguchi A, Gonzalez FJ, Sakai J, Kodama T. Transcriptome Analysis of K-877 (a Novel Selective PPARalpha Modulator (SPPARMalpha))-Regulated Genes in Primary Human Hepatocytes and the Mouse Liver. J Atheroscler Thromb. 2015;22:754–772. doi: 10.5551/jat.28720

20. Decano JL, Singh SA, Gasparotto Bueno C, Ho Lee L, Halu A, Chelvanambi S, Matamalas JT, Zhang H, Mlynarchik AK, Qiao J, et al. Systems Approach to Discovery of Therapeutic Targets for Vein Graft Disease: PPARalpha Pivotally Regulates Metabolism, Activation, and Heterogeneity of Macrophages and Lesion Development. Circulation. 2021;143:2454–2470. doi: 10.1161/CIRCULATIONAHA.119.043724

21. Iwata H, Osborn EA, Ughi GJ, Murakami K, Goettsch C, Hutcheson JD, Mauskapf A, Mattson PC, Libby P, Singh SA, et al. Highly Selective PPARalpha (Peroxisome Proliferator-Activated Receptor alpha) Agonist Pemafibrate Inhibits Stent Inflammation and Restenosis Assessed by Multimodality Molecular-Microstructural Imaging. J Am Heart Assoc. 2021;10:e020834. doi: 10.1161/JAHA.121.020834

22. Sasaki Y, Asahiyama M, Tanaka T, Yamamoto S, Murakami K, Kamiya W, Matsumura Y, Osawa T, Anai M, Fruchart JC, et al. Pemafibrate, a selective PPARalpha modulator, prevents non-alcoholic steatohepatitis development without reducing the hepatic triglyceride content. Sci Rep. 2020;10:7818. doi: 10.1038/s41598-020-64902-8

23. Fruchart JC, Santos RD, Aguilar-Salinas C, Aikawa M, Al Rasadi K, Amarenco P, Barter PJ, Ceska R, Corsini A, Despres JP, et al. The selective peroxisome proliferator-activated receptor alpha modulator (SPPARMalpha) paradigm: conceptual framework and therapeutic potential : A consensus statement from the International Atherosclerosis Society (IAS) and the Residual Risk Reduction Initiative (R3i) Foundation. Cardiovasc Diabetol. 2019;18:71. doi: 10.1186/s12933-019-0864-7

24. Pradhan AD, Glynn RJ, Fruchart JC, MacFadyen JG, Zaharris ES, Everett BM, Campbell SE, Oshima R, Amarenco P, Blom DJ, et al. Triglyceride Lowering with Pemafibrate to Reduce Cardiovascular Risk. N Engl J Med. 2022;387:1923–1934. doi: 10.1056/NEJMoa2210645

25. Honda S, Miyamoto T, Watanabe T, Narumi T, Kadowaki S, Honda Y, Otaki Y, Hasegawa H, Netsu S, Funayama A, et al. A novel mouse model of aortic valve stenosis induced by direct wire injury. Arterioscler Thromb Vasc Biol. 2014;34:270–278. doi: 10.1161/ATVBAHA.113.302610

26. Araki M, Nakagawa Y, Oishi A, Han SI, Wang Y, Kumagai K, Ohno H, Mizunoe Y, Iwasaki H, Sekiya M, et al. The Peroxisome Proliferator-Activated Receptor alpha (PPARalpha) Agonist Pemafibrate Protects against Diet-Induced Obesity in Mice. Int J Mol Sci. 2018;19. doi: 10.3390/ijms19072148

27. Egusa G, Ohno H, Nagano G, Sagawa J, Shinjo H, Yamamoto Y, Himeno N, Morita Y, Kanai A, Baba R, et al. Selective activation of PPARalpha maintains thermogenic capacity of beige adipocytes. iScience. 2023;26:107143. doi: 10.1016/j.isci.2023.107143

28. Kawakami R, Katsuki S, Travers R, Romero DC, Becker-Greene D, Passos LSA, Higashi H, Blaser MC, Sukhova GK, Buttigieg J, et al. S100A9-RAGE Axis Accelerates Formation of Macrophage-Mediated Extracellular Vesicle Microcalcification in Diabetes Mellitus. Arterioscler Thromb Vasc Biol. 2020;40:1838–1853. doi: 10.1161/ATVBAHA.118.314087

29. Averill MM, Kerkhoff C, Bornfeldt KE. S100A8 and S100A9 in cardiovascular biology and disease. Arterioscler Thromb Vasc Biol. 2012;32:223–229. doi: 10.1161/ATVBAHA.111.236927

30. Van Campenhout A, Golledge J. Osteoprotegerin, vascular calcification and atherosclerosis. Atherosclerosis. 2009;204:321-329. doi: 10.1016/j.atherosclerosis.2008.09.033

31. Gu X, Masters KS. Role of the Rho pathway in regulating valvular interstitial cell phenotype and nodule formation. Am J Physiol Heart Circ Physiol. 2011;300:H448–458. doi: 10.1152/ajpheart.01178.2009

32. Gu X, Masters KS. Role of the MAPK/ERK pathway in valvular interstitial cell calcification. Am J Physiol Heart Circ Physiol. 2009;296:H1748–1757. doi: 10.1152/ajpheart.00099.2009

33. Lee SJ, Jeong JY, Oh CJ, Park S, Kim JY, Kim HJ, Doo Kim N, Choi YK, Do JY, Go Y, et al. Pyruvate Dehydrogenase Kinase 4 Promotes Vascular Calcification via SMAD1/5/8 Phosphorylation. Sci Rep. 2015;5:16577. doi: 10.1038/srep16577

34. Cai Y, Wang XL, Flores AM, Lin T, Guzman RJ. Inhibition of endo-lysosomal function exacerbates vascular calcification. Sci Rep. 2018;8:3377. doi: 10.1038/s41598-017-17540-6

35. Sanchez-Bayuela T, Peral-Rodrigo M, Parra-Izquierdo I, Lopez J, Gomez C, Montero O, Perez-Riesgo E, San Roman JA, Butcher JT, Sanchez Crespo M, et al. Inflammation via JAK-STAT/HIF-1alpha Drives Metabolic Changes in Pentose Phosphate Pathway and Glycolysis That Support Aortic Valve Cell Calcification. Arterioscler Thromb Vasc Biol. 2025;45:e232–e249. doi: 10.1161/ATVBAHA.124.322375

36. Zhao Y, Sun Z, Li L, Yuan W, Wang Z. Role of Collagen in Vascular Calcification. J Cardiovasc Pharmacol. 2022;80:769–778. doi: 10.1097/FJC.0000000000001359

37. Anousakis-Vlachochristou N, Athanasiadou D, Carneiro KMM, Toutouzas K. Focusing on the Native Matrix Proteins in Calcific Aortic Valve Stenosis. JACC Basic Transl Sci. 2023;8:1028–1039. doi: 10.1016/j.jacbts.2023.01.009

38. Barth M, Selig JI, Klose S, Schomakers A, Kiene LS, Raschke S, Boeken U, Akhyari P, Fischer JW, Lichtenberg A. Degenerative aortic valve disease and diabetes: Implications for a link between proteoglycans and diabetic disorders in the aortic valve. Diab Vasc Dis Res. 2019;16:254–269. doi: 10.1177/1479164118817922

39. Gollmann-Tepekoylu C, Graber M, Hirsch J, Mair S, Naschberger A, Polzl L, Nagele F, Kirchmair E, Degenhart G, Demetz E, et al. Toll-Like Receptor 3 Mediates Aortic Stenosis Through a Conserved Mechanism of Calcification. Circulation. 2023;147:1518–1533. doi: 10.1161/CIRCULATIONAHA.122.063481

40. Cho DI, Kang HJ, Jeon JH, Eom GH, Cho HH, Kim MR, Cho M, Jeong HY, Cho HC, Hong MH, et al. Antiinflammatory activity of ANGPTL4 facilitates macrophage polarization to induce cardiac repair. JCI Insight. 2019;4. doi: 10.1172/jci.insight.125437

41. Singh P, Dejager L, Amand M, Theatre E, Vandereyken M, Zurashvili T, Singh M, Mack M, Timmermans S, Musumeci L, et al. DUSP3 Genetic Deletion Confers M2-like Macrophage-Dependent Tolerance to Septic Shock. J Immunol. 2015;194:4951–4962. doi: 10.4049/jimmunol.1402431

42. Proudfoot D, Shanahan CM. Molecular mechanisms mediating vascular calcification: role of matrix Gla protein. Nephrology (Carlton). 2006;11:455–461. doi: 10.1111/j.1440-1797.2006.00660.x

43. Liu XH, Zhou JT, Yan CX, Cheng C, Fan JN, Xu J, Zheng Q, Bai Q, Li Z, Li S, et al. Single-cell RNA sequencing reveals a novel inhibitory effect of ApoA4 on NAFL mediated by liver-specific subsets of myeloid cells. Front Immunol. 2022;13:1038401. doi: 10.3389/fimmu.2022.1038401

44. Troutt JS, Alborn WE, Cao G, Konrad RJ. Fenofibrate treatment increases human serum proprotein convertase subtilisin kexin type 9 levels. J Lipid Res. 2010;51:345–351. doi: 10.1194/jlr.M000620

45. Sahebkar A. Circulating levels of proprotein convertase subtilisin kexin type 9 are elevated by fibrate therapy: a systematic review and meta-analysis of clinical trials. Cardiol Rev. 2014;22:306–312. doi: 10.1097/CRD.0000000000000025

46. Mayne J, Dewpura T, Raymond A, Cousins M, Chaplin A, Lahey KA, Lahaye SA, Mbikay M, Ooi TC, Chretien M. Plasma PCSK9 levels are significantly modified by statins and fibrates in humans. Lipids Health Dis. 2008;7:22. doi: 10.1186/1476-511X-7-22

47. Blaser MC, Buffolo F, Halu A, Turner ME, Schlotter F, Higashi H, Pantano L, Clift CL, Saddic LA, Atkins SK, et al. Multiomics of Tissue Extracellular Vesicles Identifies Unique Modulators of Atherosclerosis and Calcific Aortic Valve Stenosis. Circulation. 2023;148:661–678. doi: 10.1161/CIRCULATIONAHA.122.063402

48. Monga S, Valkovic L, Myerson SG, Neubauer S, Rider OJ, Mahmod M. Fenofibrate modulates cardiac energy metabolism in moderate-severe aortic stenosis: a mechanistic double-blinded randomised controlled MR trial. European Heart Journal. 2024;45. doi: 10.1093/eurheartj/ehae666.255

49. Kaltoft M, Langsted A, Nordestgaard BG. Triglycerides and remnant cholesterol associated with risk of aortic valve stenosis: Mendelian randomization in the Copenhagen General Population Study. Eur Heart J. 2020;41:2288–2299. doi: 10.1093/eurheartj/ehaa172

50. Varga A, Hegele RA. Triglyceride-rich particles: new actors in valvular aortic stenosis. Eur Heart J. 2020;41:2300–2303. doi: 10.1093/eurheartj/ehaa416

51. Chen Z, Liu L, Jiao X, Zhang Y, Wang F, Chen Y, Lan Z, Liu X. The association between triglyceride to high-density-lipoprotein cholesterol ratio and calcific aortic valve disease: a retrospective study. BMC Cardiovasc Disord. 2024;24:708. doi: 10.1186/s12872-024-04372-2

52. Turner ME, Aikawa E. Updating the paradigm: inflammation as a targetable modulator of medial vascular calcification. Cardiovasc Res. 2023;119:2259–2261. doi: 10.1093/cvr/cvad139

53. Bartoli-Leonard F, Zimmer J, Aikawa E. Innate and adaptive immunity: the understudied driving force of heart valve disease. Cardiovasc Res. 2021;117:2506–2524. doi: 10.1093/cvr/cvab273

54. Niepmann ST, Steffen E, Zietzer A, Adam M, Nordsiek J, Gyamfi-Poku I, Piayda K, Sinning JM, Baldus S, Kelm M, et al. Graded murine wire-induced aortic valve stenosis model mimics human functional and morphological disease phenotype. Clin Res Cardiol. 2019;108:847–856. doi: 10.1007/s00392-019-01413-1

55. Lu J, Xie S, Deng Y, Xie X, Liu Y. Blocking the NLRP3 inflammasome reduces osteogenic calcification and M1 macrophage polarization in a mouse model of calcified aortic valve stenosis. Atherosclerosis. 2022;347:28–38. doi: 10.1016/j.atherosclerosis.2022.03.005

56. Ji YY, Liu JT, Liu N, Wang ZD, Liu CH. PPARalpha activator fenofibrate modulates angiotensin II-induced inflammatory responses in vascular smooth muscle cells via the TLR4-dependent signaling pathway. Biochem Pharmacol. 2009;78:1186–1197. doi: 10.1016/j.bcp.2009.06.095

57. Kooistra T, Verschuren L, de Vries-van der Weij J, Koenig W, Toet K, Princen HM, Kleemann R. Fenofibrate reduces atherogenesis in ApoE*3Leiden mice: evidence for multiple antiatherogenic effects besides lowering plasma cholesterol. Arterioscler Thromb Vasc Biol. 2006;26:2322–2330. doi: 10.1161/01.ATV.0000238348.05028.14

58. Zandbergen F, Plutzky J. PPARalpha in atherosclerosis and inflammation. Biochim Biophys Acta. 2007;1771:972-982. doi: 10.1016/j.bbalip.2007.04.021

59. Goody PR, Hosen MR, Christmann D, Niepmann ST, Zietzer A, Adam M, Bonner F, Zimmer S, Nickenig G, Jansen F. Aortic Valve Stenosis: From Basic Mechanisms to Novel Therapeutic Targets. Arterioscler Thromb Vasc Biol. 2020;40:885–900. doi: 10.1161/ATVBAHA.119.313067

60. 60. Report on the Deliberation Results, Parmodia Tab. 0.1 mg, Pemafibrate (JAN), Kowa Comapny, Ltd. Pharmaceuticals and Medical Devices Agency (PMDA). 2017.

61. Goto S, Rogers MA, Blaser MC, Higashi H, Lee LH, Schlotter F, Body SC, Aikawa M, Singh SA, Aikawa E. Standardization of Human Calcific Aortic Valve Disease in vitro Modeling Reveals Passage-Dependent Calcification. Front Cardiovasc Med. 2019;6:49. doi: 10.3389/fcvm.2019.00049

62. Hennuyer N, Duplan I, Paquet C, Vanhoutte J, Woitrain E, Touche V, Colin S, Vallez E, Lestavel S, Lefebvre P, et al. The novel selective PPARalpha modulator (SPPARMalpha) pemafibrate improves dyslipidemia, enhances reverse cholesterol transport and decreases inflammation and atherosclerosis. Atherosclerosis. 2016;249:200–208. doi: 10.1016/j.atherosclerosis.2016.03.003

63. Clift CL, Blaser MC, Bartoli-Leonard F, Schlotter F, Higashi H, Kuraoka S, Kasai T, Turner ME, Pham T, Perez KA, et al. Sexual Dimorphism of Plasma and Tissue Proteomes in Human Calcific Aortic Valve Stenosis Pathogenesis-Brief Report. Arterioscler Thromb Vasc Biol. 2025;45:1928–1934. doi: 10.1161/ATVBAHA.125.322560

64. Tyanova S, Temu T, Sinitcyn P, Carlson A, Hein MY, Geiger T, Mann M, Cox J. The Perseus computational platform for comprehensive analysis of (prote)omics data. Nat Methods. 2016;13:731–740. doi: 10.1038/nmeth.3901

65. Hulsen T, de Vlieg J, Alkema W. BioVenn - a web application for the comparison and visualization of biological lists using area-proportional Venn diagrams. BMC Genomics. 2008;9:488. doi: 10.1186/1471-2164-9-488

66. Szklarczyk D, Kirsch R, Koutrouli M, Nastou K, Mehryary F, Hachilif R, Gable AL, Fang T, Doncheva NT, Pyysalo S, et al. The STRING database in 2023: protein-protein association networks and functional enrichment analyses for any sequenced genome of interest. Nucleic Acids Res. 2023;51:D638–D646. doi: 10.1093/nar/gkac1000

67. Evangelista JE, Xie Z, Marino GB, Nguyen N, Clarke DJB, Ma’ayan A. Enrichr-KG: bridging enrichment analysis across multiple libraries. Nucleic Acids Res. 2023;51:W168–W179. doi: 10.1093/nar/gkad393

