## Supplemental Figures for "Selective PPAR-α activation with pemafibrate attenuates macrophage-mediated progression of calcific aortic valve disease"

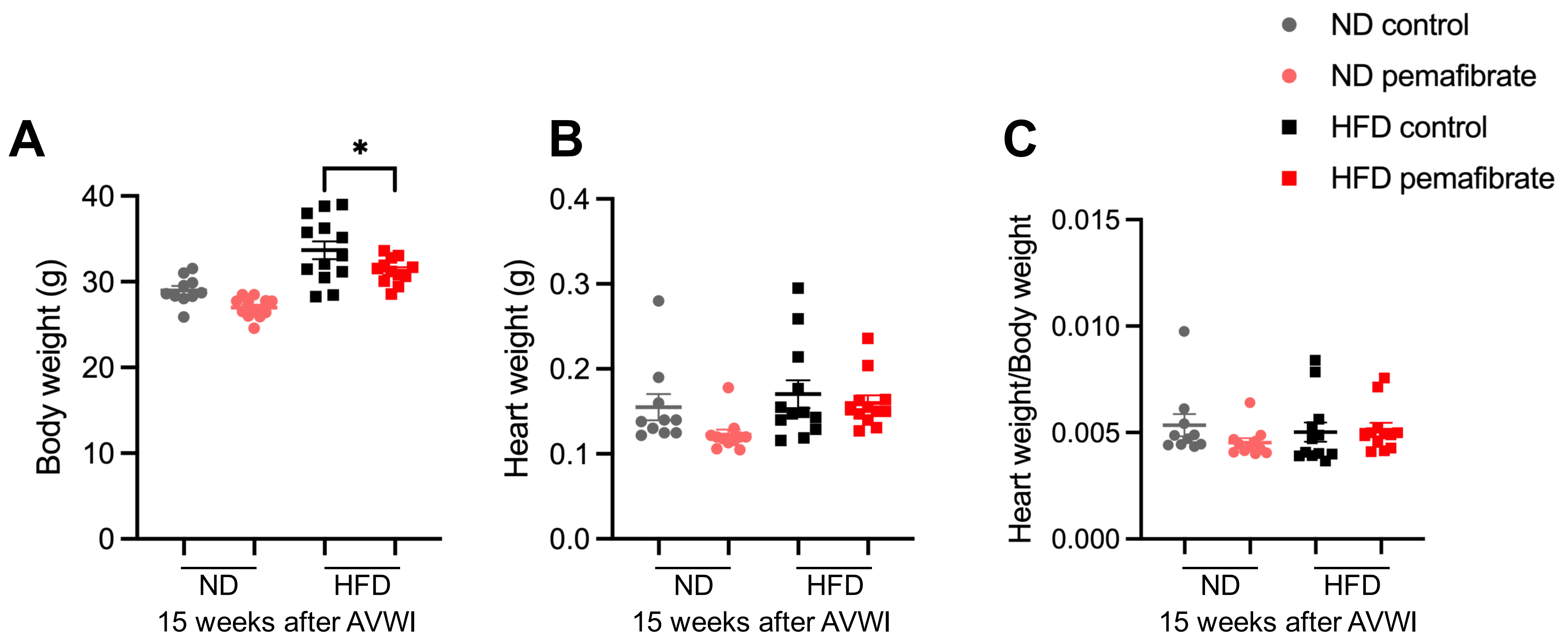


**Supplementary Figure S1: Body weight and heart weight in AVWI mice with pemafibrate treatment at week 15**
**(A)** Body weight (g), **(B)** Heart weight (g), and **(C)** Relative values of heart weight/body weight in AVWI model at 15 weeks after surgery. Mean±SEM. N=10-12/group. Ordinary two-way ANOVA followed by Bonferroni *post hoc* test performed for statistical analysis between control and pemafibrate in ND and HFD. *p<0.05.

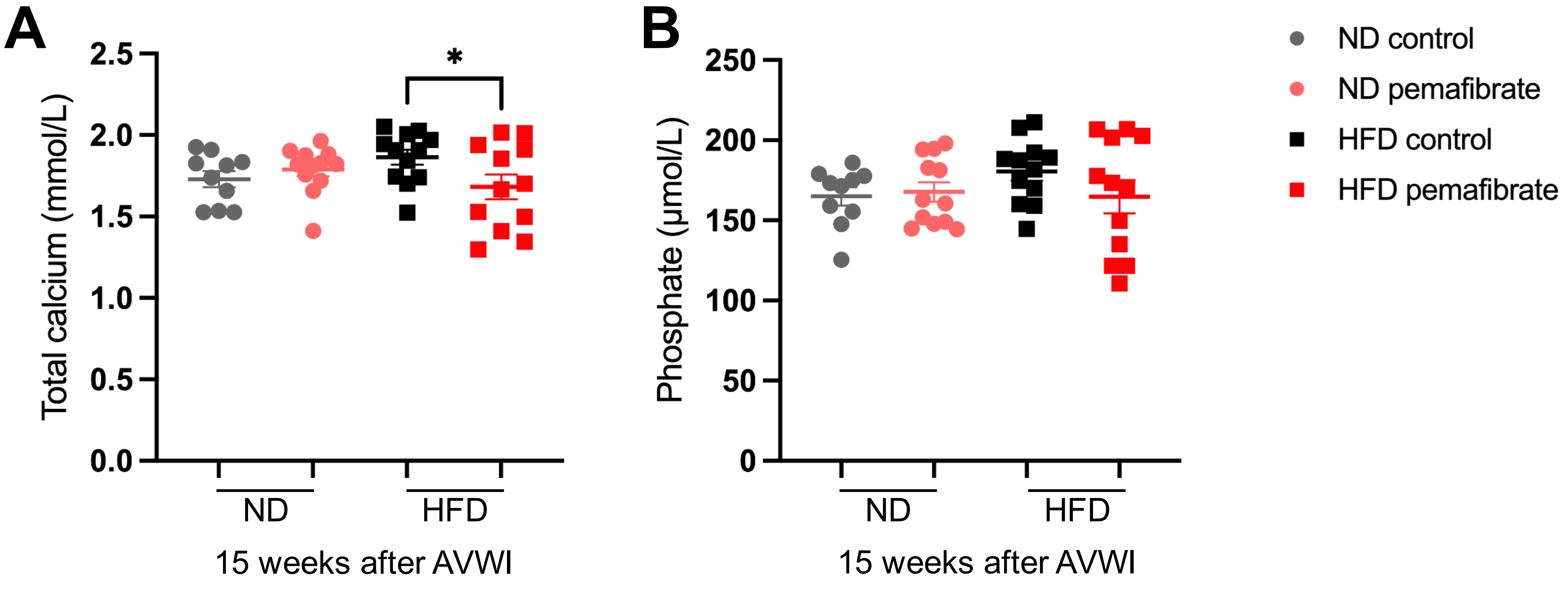


**Supplementary Figure S2: Plasma calcium decreased and phosphate level did not change with pemafibrate in AVWI mice at week 15**

**(A)** Total calcium (mmol/L) and **(B)** phosphate (µmol/L) in plasma in AVWI model at 15 weeks after surgery. Mean±SEM. N=10-12/group. Ordinary two-way ANOVA followed by Bonferroni *post hoc* test performed for statistical analysis between control and pemafibrate in ND and HFD. *p<0.05.

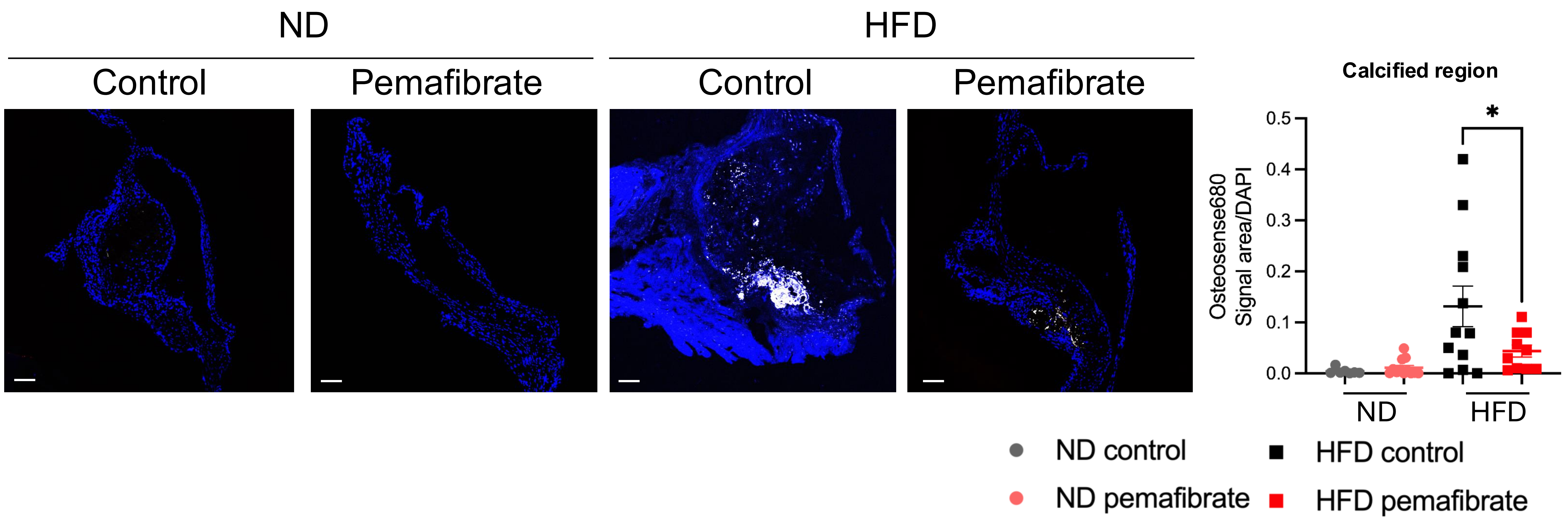


**Supplementary Figure S3: Pemafibrate suppressed aortic valve leaftlet calcification in HFD group**
Osteosense680 signal area in the aortic valve 15 weeks after AVWI (Blue: DAPI, White: OsteoSense680). Mean±SEM. N=10-12/group. Scale bar = 100μm. Ordinary two-way ANOVA followed by Bonferroni *post hoc* test performed for statistical analysis between control and pemafibrate in ND and HFD. *p<0.05.

### Correlation with TG and echo parameters (15 weeks) in HFD group

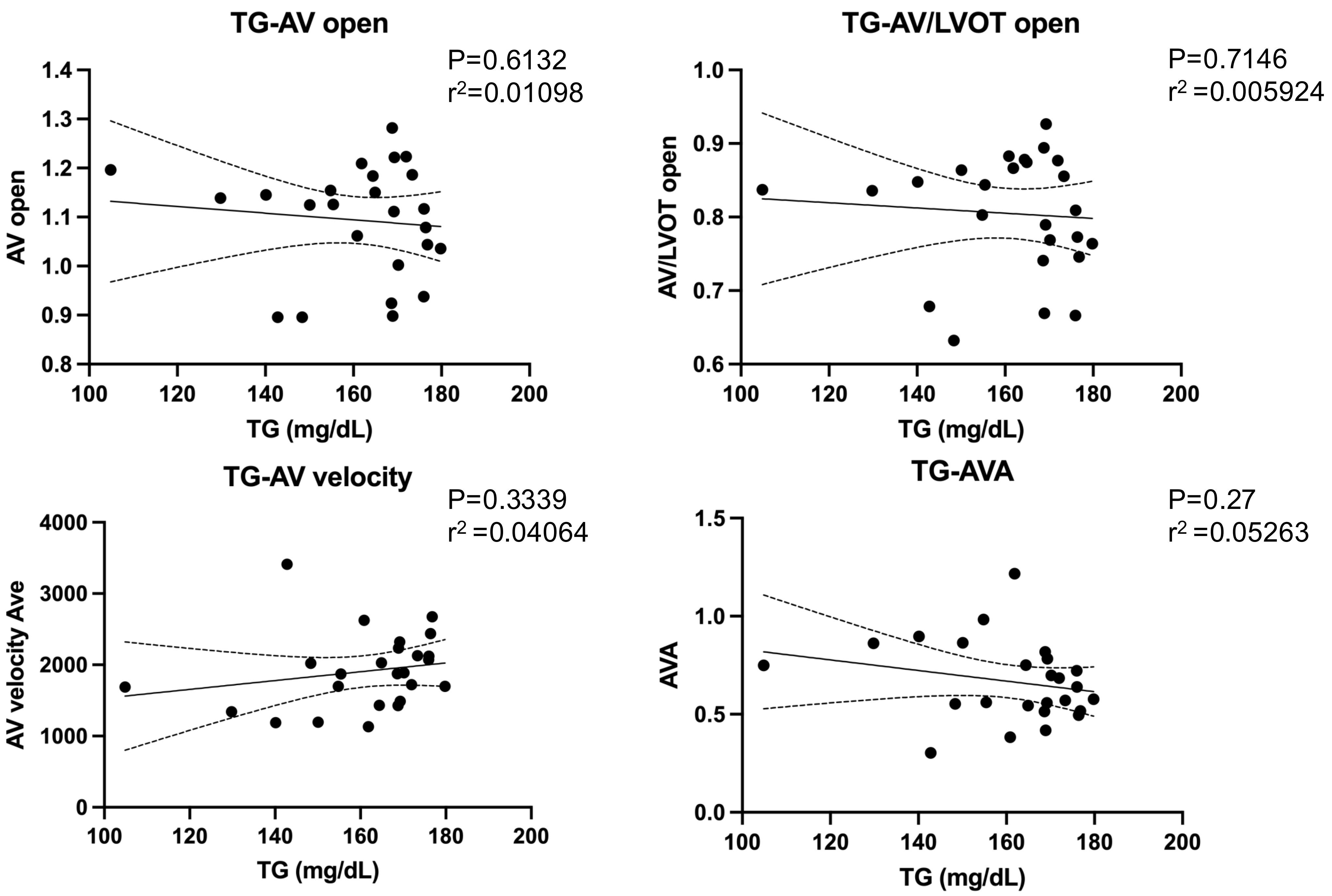


**Supplementary Figure S4: Echocardiography parameters did not correlate with plasma triglyceride level**
Plasma triglyceride levels plotted against echocardiography parameters. The parameters were acquired at the 15-week time point in high-fat diet-fed group. Linear regression plotted with 95% confidence bands of the best-fit line.

### Correlation with TG and histology parameters (15 weeks) in HFD group

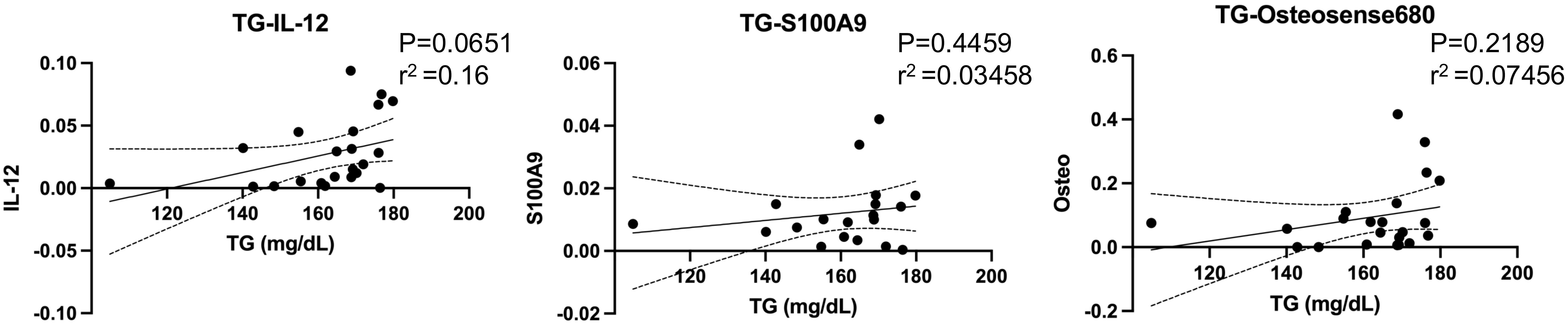


**Supplementary Figure S5: Plasma triglyceride levels did not correlate with aortic valve inflammation markers or calcified aortic valve area using histopathology**

Plasma triglyceride levels plotted against inflammation markers (IL-12 and S100A9) or calcification (Osteosense) in the aortic valve leaflet. The parameters were acquired at the 15-week time point in high-fat diet-fed group. Linear regression plotted with 95% confidence bands of the best-fit line.

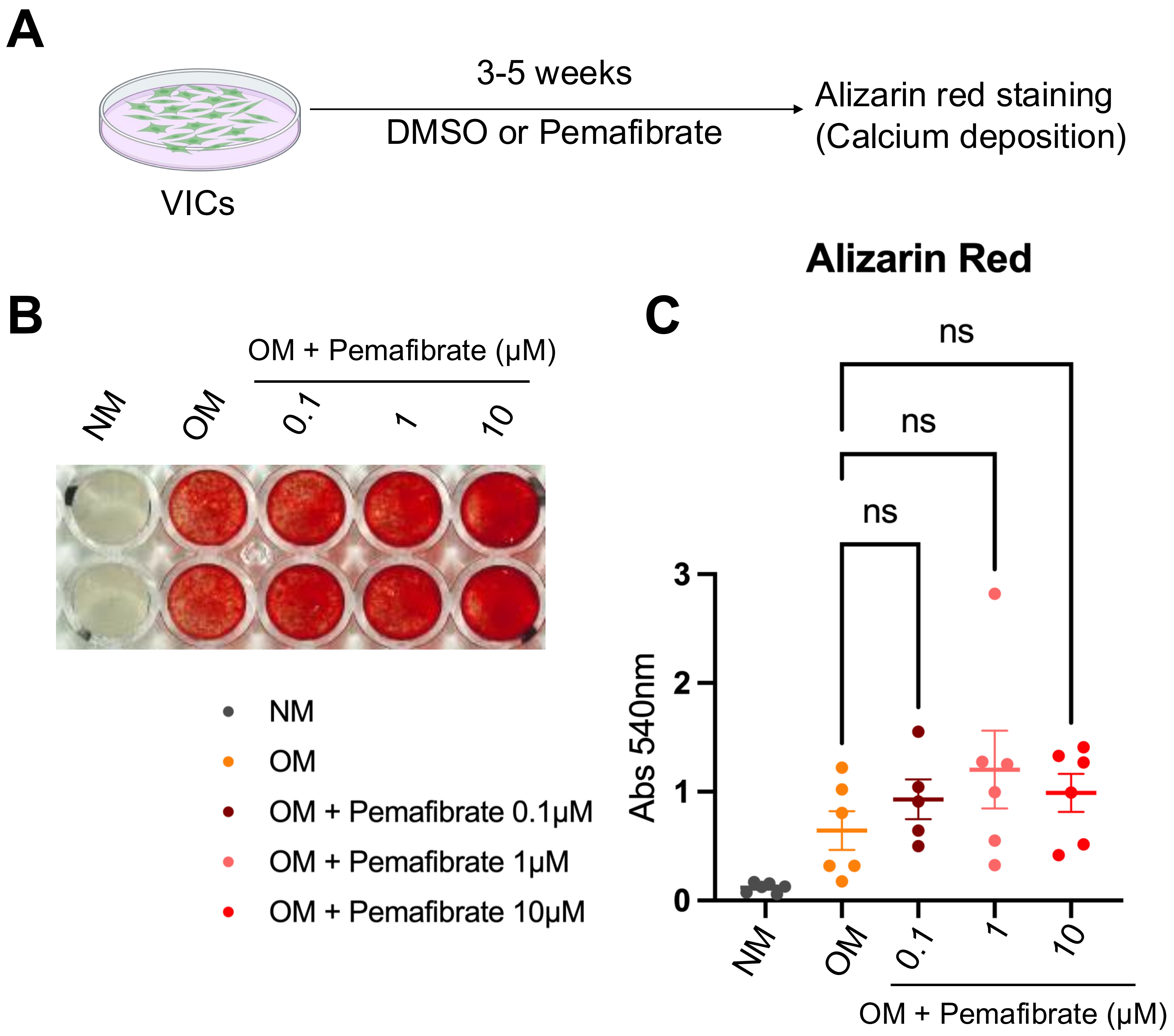


**Supplementary Figure S6: Pemafibrate did not directly inhibit calcification of human primary VICs**
**(A)** Schematic of *in vitro* assay in VICs treated with pemafibrate. **(B)** Representative images of alizarin red staining in VICs (NM, normal media; OM, osteogenic media). **(C)** Donor-specific quantification of alizarin red staining in human VICs. The plots represented the values from independent duplicate wells. Ordinary one-way ANOVA followed by Dunnett's multiple comparison test.

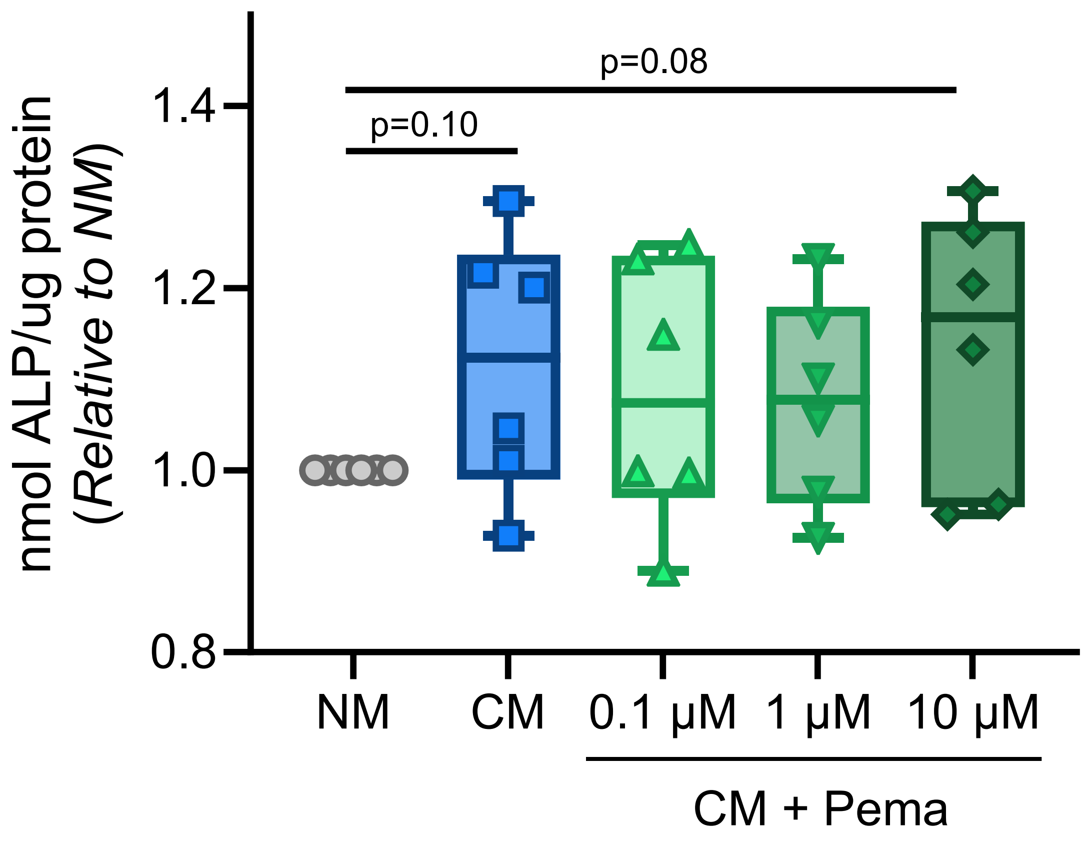


**Supplementary Figure S7: Conditioned media from THP-1 macrophage-like cells did not change ALP activity in VICs, regardless of pemafibrate treatment**

ALP activity value was normalized by protein amount. N=6 donors per group.

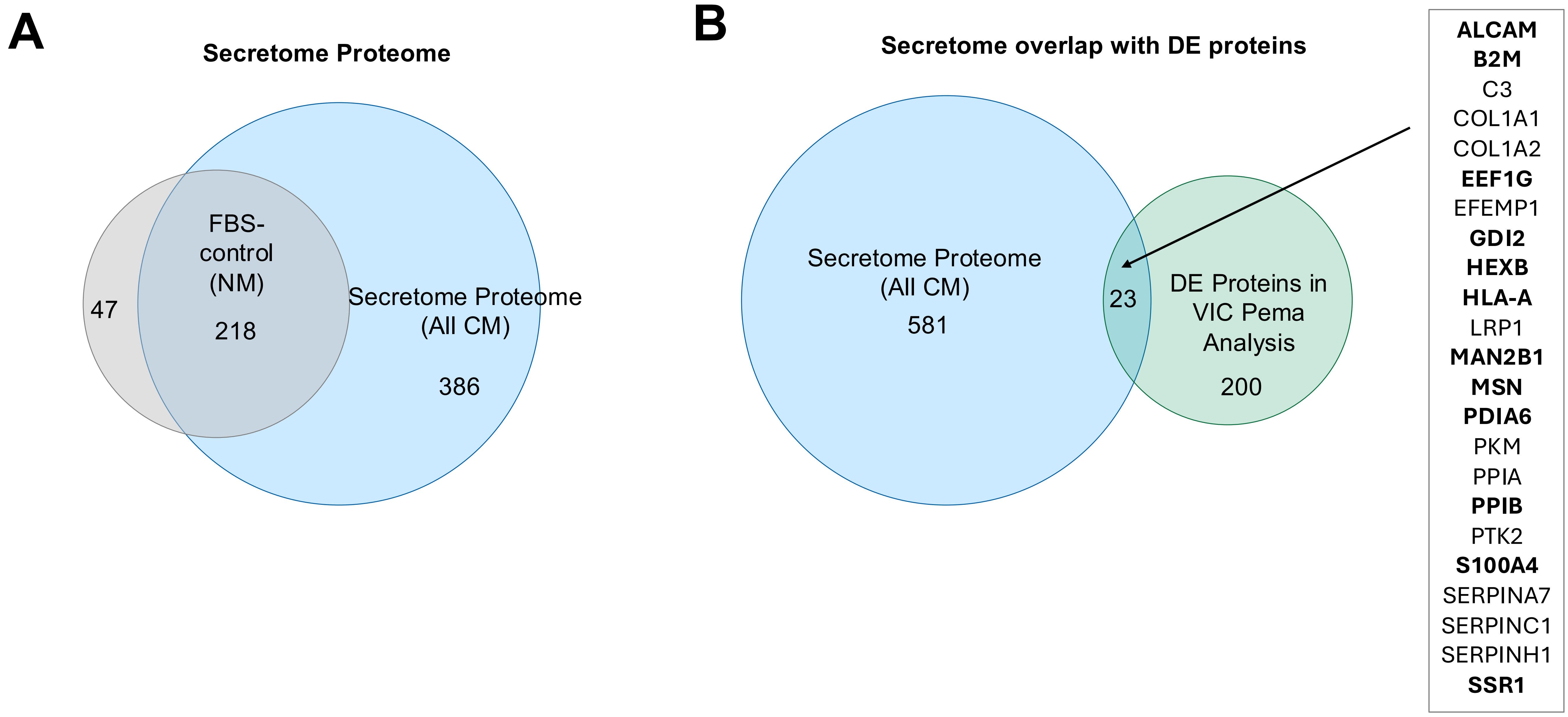


**Supplementary Figure S8: 5% of differentially enriched proteins in the VIC proteomic analysis proteins are also identified in the secretome and reflect proteins that may in-part or wholly be derived from THP-1 cells.**
**(A)** Overlap of protein IDs found in THP-1 conditioned media (secretome proteome, n=604) and the experimental NM control (n=265), containing only fetal bovine serum and media. The unique secretome proteome (n=386, Supplementary Table S10) **(B)** Overlap of protein IDs from the secretome compared to all differentially enriched proteins in the pemafibrate analysis presented in Figure 5 (n=223). 23 overlap with the entire secretome including FBS-derived and 11 (bolded in in Figure) are only identified in CM. Experimental VICs were treated with pooled conditioned media, thus n=1 for each condition (NM, CM, CM + Pema 0.1, 1, 10 μM)

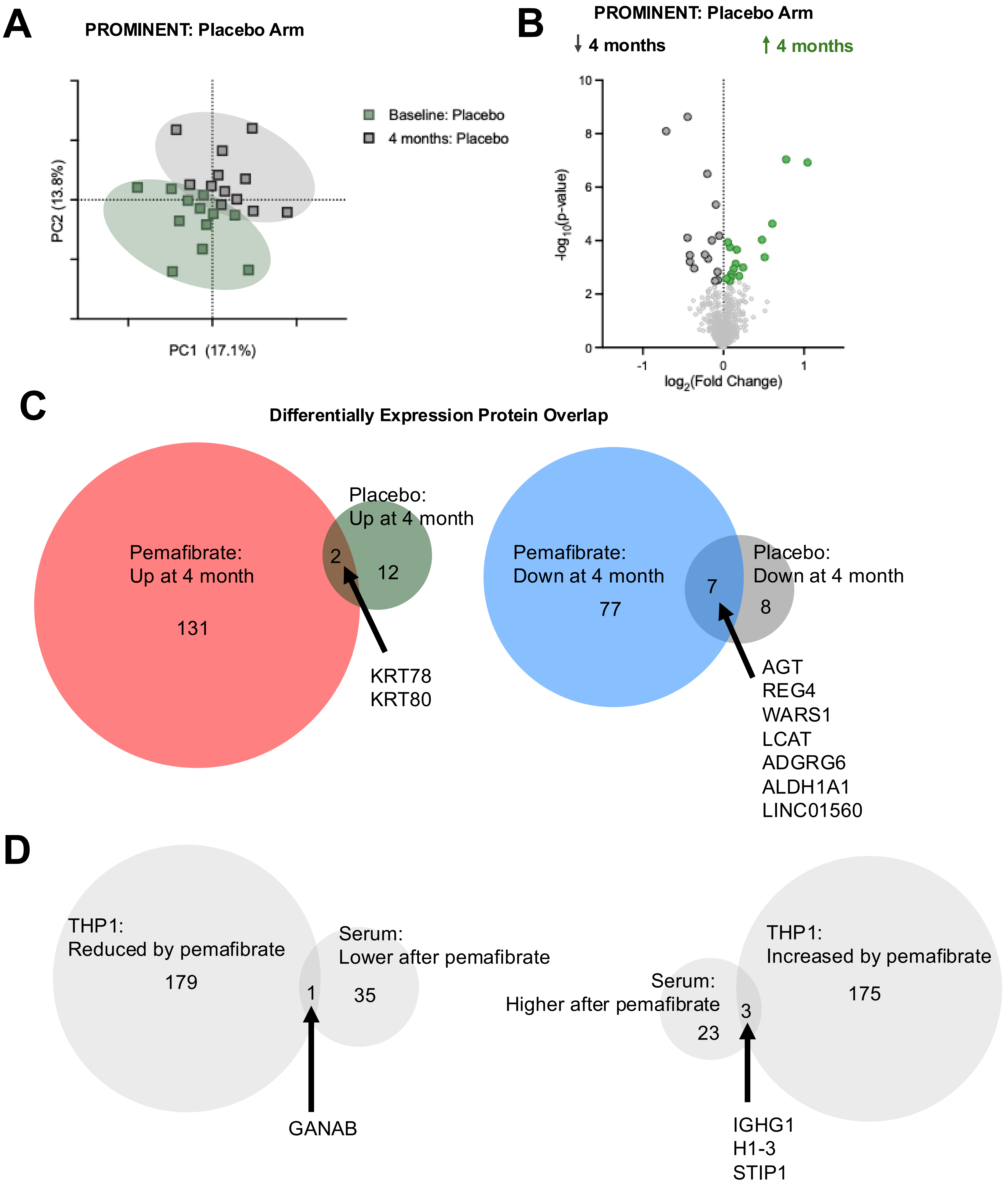


**Supplementary Figure S9: Serum proteomic analysis of placebo group from PROMINENT**
**(A)** Principal component analysis serum proteome (684 IDs) of patients from the placebo arm at baseline and following 4 months. Effect of participant variability removed using a generalized linear model. **(B)** Differential enrichment analysis comparing baseline to 4 months following pemafibrate using a T-test. Highlighted in red and blue and red are proteins with FDR q<0.05. **(C)** Overlap of differentially enriched proteins in placebo and pemafibrate arm. **(D)** Z-score normalized protein abundance in placebo arm plotted against the pemafibrate arm of the differentially enriched proteins in the pemafibrate-arm (Figure 6E). Subsets of proteins selected for further assessment with large change in pemafibrate Z-score normalized abundance (>0.6) and small change in the placebo arm (<0.1).

**Supplemental Table S1. Taqman qPCR probe list**

| Gene name | Taqman probe ID |
|---|---|
| *MSX2* | Hs00751239_s1 |
| *RUNX2* | Hs00231692_m1 |
| *OPN* | Hs00959010_m1 |
| *TLR4* | Hs00152939_m1 |
| *ICAM1* | Hs99999152_m1 |
| *IL1B* | Hs00174097_m1 |
| *IL6* | Hs00985639_m1 |
| *IL12* | Hs00233688_m1 |
| *S100A8* | Hs00374264_g1 |
| *S100A9* | Hs00610058_m1 |
| *MMP1* | Hs0089660_g1 |
| *MMP2* | Hs01548724_m3 |
| *MMP9* | Hs00957555_m1 |
| *THBS1* | Hs00962908_m1 |
| *HIF1A* | Hs00153153_m1 |
| *WARS* | Hs00188259 |
| *HMOX1* | Hs01110250_m1 |
| *XBP1* | Hs00231936_m1 |
| *ENO1* | Hs00361415_m1 |
| *NAMPT* | Hs00237184_m1 |
| *CCL2* | Hs00234140_m1 |
| *CCL4* | Hs00234142_g1 |
| *CCL5* | Hs00982282_m1 |
| *CCL8* | Hs04187715_m1 |
| *CXCL1* | Hs00236937_m1 |
| *CXCL6* | Hs00237017_m1 |
| *CXCL8* | Hs00174103_m1 |
| *CXCL10* | Hs01124251_g1 |
| *CXCL11* | Hs04187682_g1 |
| *CXCL12* | Hs03676656_mH |
| *CXCR4* | Hs00607978_s1 |

**Supplemental Table S2. Echocardiography analysis in aortic valve wire injury (AVWI) model with/without pemafibrate treatment**

| | | ND | | | HFD | | |
|---|---|---|---|---|---|---|---|
| **Parameter** | **Time** | **Control n=9** | **Pemafibrate n=12** | **p-value** | **Control n=11** | **Pemafibrate n=12** | **p-value** |
| *Aortic valve parameters* | | | | | | | |
| AV open (mm) | Pre | 1.138 ± 0.027 | 1.161 ± 0.017 | 0.486 | 1.188 ± 0.033 | 1.213 ± 0.017 | 0.507 |
| | 8 weeks | 1.044 ± 0.033 | 1.122 ± 0.019 | **0.057** | 1.020 ± 0.027 | 1.154 ± 0.020 | **<0.001** |
| | 15 weeks | 1.010 ± 0.016 | 1.102 ± 0.023 | **0.004** | 1.002 ± 0.026 | 1.156 ± 0.021 | **<0.001** |
| LVOT open (mm) | Pre | 1.340 ± 0.017 | 1.313 ± 0.028 | 0.416 | 1.382 ± 0.027 | 1.377 ± 0.014 | 0.868 |
| | 8 weeks | 1.337 ± 0.030 | 1.323 ± 0.016 | 0.686 | 1.368 ± 0.018 | 1.368 ± 0.018 | 0.996 |
| | 15 weeks | 1.359 ± 0.019 | 1.383 ± 0.026 | 0.477 | 1.354 ± 0.015 | 1.360 ± 0.021 | 0.803 |
| AV open/LVOT open | Pre | 0.852 ± 0.018 | 0.887 ± 0.017 | 0.166 | 0.858 ± 0.013 | 0.881 ± 0.009 | 0.181 |
| | 8 weeks | 0.781 ± 0.016 | 0.849 ± 0.015 | **0.005** | 0.746 ± 0.018 | 0.844 ± 0.010 | **<0.001** |
| | 15 weeks | 0.744 ± 0.014 | 0.798 ± 0.014 | **0.013** | 0.740 ± 0.019 | 0.850 ± 0.009 | **<0.001** |
| AVA ($cm^2$) | Pre | 0.674 ± 0.050 | 0.711 ± 0.055 | 0.621 | 1.030 ± 0.061 | 0.983 ± 0.067 | 0.611 |
| | 8 weeks | 0.543 ± 0.040 | 0.741 ± 0.104 | 0.1 | 0.538 ± 0.040 | 0.795 ± 0.049 | **<0.001** |
| | 15 weeks | 0.607 ± 0.049 | 0.765 ± 0.064 | **0.065** | 0.531 ± 0.033 | 0.725 ± 0.052 | **0.005** |
| AV velocity (m/s) | Pre | 1361.463 ± 66.186 | 1300.603 ± 46.664 | 0.463 | 1147.114 ± 29.232 | 1205.540 ± 37.992 | 0.236 |
| | 8 weeks | 1952.833 ± 162.221 | 1485.379 ± 114.970 | **0.03** | 2351.942 ± 179.072 | 1634.373 ± 81.305 | **0.002** |
| | 15 weeks | 1996.764 ± 105.825 | 1402.631 ± 69.087 | **<0.001** | 2251.698 ± 122.612 | 1679.140 ± 112.238 | **0.002** |
| LVOT velocity (m/s) | Pre | 620.584 ± 25.236 | 668.720 ± 30.173 | 0.235 | 781.687 ± 33.384 | 787.503 ± 44.226 | 0.917 |
| | 8 weeks | 778.419 ± 82.235 | 703.125 ± 51.591 | 0.45 | 828.895 ± 48.415 | 865.415 ± 37.016 | 0.556 |
| | 15 weeks | 823.826 ± 65.812 | 686.460 ± 15.703 | **0.07** | 805.408 ± 30.339 | 800.596 ± 24.710 | 0.903 |
| AV velocity/LVOT velocity | Pre | 2.227 ± 0.133 | 1.986 ± 0.108 | 0.176 | 1.495 ± 0.064 | 1.594 ± 0.108 | 0.44 |
| | 8 weeks | 2.757 ± 0.296 | 2.232 ± 0.312 | 0.237 | 2.903 ± 0.233 | 1.919 ± 0.110 | **0.002** |
| | 15 weeks | 2.515 ± 0.194 | 2.058 ± 0.117 | 0.063 | 2.837 ± 0.211 | 2.103 ± 0.127 | **0.008** |
| *Cardiac parameters* | | | | | | | |
| EF (%) | Pre | 49.763 ± 2.069 | 51.155 ± 5.154 | 0.814 | 48.495 ± 1.376 | 49.159 ± 1.332 | 0.732 |
| | 8 weeks | 60.078 ± 3.822 | 53.481 ± 2.406 | 0.163 | 52.583 ± 2.368 | 53.718 ± 3.359 | 0.785 |
| | 15 weeks | 52.282 ± 2.328 | 53.239 ± 2.974 | 0.803 | 47.039 ± 2.565 | 49.082 ± 1.947 | 0.532 |
| FS (%) | Pre | 24.980 ± 1.220 | 25.953 ± 3.224 | 0.792 | 24.254 ± 0.815 | 24.639 ± 0.812 | 0.741 |
| | 8 weeks | 32.276 ± 2.757 | 27.400 ± 1.540 | 0.142 | 27.086 ± 1.506 | 28.001 ± 2.309 | 0.744 |
| | 15 weeks | 26.634 ± 1.415 | 27.605 ± 2.127 | 0.708 | 23.786 ± 1.482 | 24.829 ± 1.120 | 0.58 |
| IVS;d (mm) | Pre | 0.595 ± 0.036 | 0.771 ± 0.038 | **0.009** | 0.597 ± 0.032 | 0.591 ± 0.024 | 0.899 |
| | 8 weeks | 0.727 ± 0.060 | 0.594 ± 0.024 | **0.061** | 0.726 ± 0.052 | 0.664 ± 0.039 | 0.344 |
| | 15 weeks | 0.689 ± 0.038 | 0.619 ± 0.037 | 0.204 | 0.698 ± 0.043 | 0.637 ± 0.035 | 0.28 |
| IVS;s (mm) | Pre | 0.950 ± 0.035 | 1.127 ± 0.072 | 0.083 | 0.972 ± 0.040 | 0.981 ± 0.027 | 0.858 |
| | 8 weeks | 1.174 ± 0.073 | 0.939 ± 0.038 | **0.012** | 1.160 ± 0.067 | 1.113 ± 0.076 | 0.65 |
| | 15 weeks | 1.062 ± 0.058 | 1.081 ± 0.072 | 0.839 | 1.167 ± 0.058 | 1.033 ± 0.037 | 0.066 |
| LV Mass ($g/m^2$, Corrected) | Pre | 72.845 ± 5.897 | 80.607 ± 3.134 | 0.27 | 72.955 ± 2.846 | 73.776 ± 2.985 | 0.844 |
| | 8 weeks | 77.001 ± 6.674 | 67.059 ± 5.764 | 0.273 | 114.842 ± 13.297 | 91.949 ± 7.812 | 0.155 |
| | 15 weeks | 95.462 ± 15.649 | 79.418 ± 6.924 | 0.366 | 129.355 ± 19.334 | 99.370 ± 8.928 | 0.177 |
| LV Vol;d (mL) | Pre | 73.273 ± 4.699 | 68.172 ± 6.161 | 0.532 | 79.642 ± 3.380 | 77.569 ± 2.569 | 0.63 |
| | 8 weeks | 70.404 ± 8.536 | 72.764 ± 6.179 | 0.825 | 101.693 ± 13.467 | 82.673 ± 6.277 | 0.219 |
| | 15 weeks | 85.446 ± 19.333 | 80.890 ± 4.304 | 0.823 | 125.622 ± 19.339 | 99.430 ± 7.746 | 0.227 |
| LV Vol;s (mL) | Pre | 37.450 ± 3.722 | 33.842 ± 5.862 | 0.623 | 41.281 ± 2.516 | 39.586 ± 1.947 | 0.599 |
| | 8 weeks | 29.971 ± 5.533 | 34.609 ± 4.059 | 0.508 | 50.261 ± 8.734 | 39.405 ± 5.027 | 0.296 |
| | 15 weeks | 43.019 ± 12.103 | 38.143 ± 3.135 | 0.705 | 71.170 ± 14.152 | 52.025 ± 6.006 | 0.231 |
| LVID;d (mm) | Pre | 4.063 ± 0.110 | 3.944 ± 0.156 | 0.556 | 4.213 ± 0.077 | 4.171 ± 0.059 | 0.665 |
| | 8 weeks | 3.955 ± 0.184 | 4.036 ± 0.138 | 0.73 | 4.591 ± 0.255 | 4.267 ± 0.123 | 0.268 |
| | 15 weeks | 4.213 ± 0.322 | 4.239 ± 0.090 | 0.941 | 4.982 ± 0.320 | 4.608 ± 0.152 | 0.305 |
| LVID;s (mm) | Pre | 3.057 ± 0.126 | 2.928 ± 0.213 | 0.624 | 3.195 ± 0.080 | 3.145 ± 0.064 | 0.632 |
| | 8 weeks | 2.710 ± 0.208 | 2.941 ± 0.141 | 0.369 | 3.368 ± 0.233 | 3.088 ± 0.162 | 0.336 |
| | 15 weeks | 3.112 ± 0.284 | 3.074 ± 0.119 | 0.906 | 3.839 ± 0.311 | 3.479 ± 0.162 | 0.317 |
| LVPW;d (mm) | Pre | 0.697 ± 0.069 | 0.686 ± 0.084 | 0.926 | 0.636 ± 0.027 | 0.667 ± 0.038 | 0.523 |
| | 8 weeks | 0.669 ± 0.046 | 0.613 ± 0.041 | 0.376 | 0.811 ± 0.055 | 0.772 ± 0.054 | 0.619 |
| | 15 weeks | 0.783 ± 0.035 | 0.670 ± 0.047 | 0.071 | 0.754 ± 0.038 | 0.728 ± 0.028 | 0.591 |
| LVPW;s (mm) | Pre | 1.081 ± 0.056 | 1.043 ± 0.090 | 0.727 | 0.987 ± 0.024 | 1.008 ± 0.038 | 0.65 |
| | 8 weeks | 1.094 ± 0.080 | 1.007 ± 0.064 | 0.405 | 1.224 ± 0.052 | 1.181 ± 0.069 | 0.623 |
| | 15 weeks | 1.200 ± 0.039 | 1.065 ± 0.059 | 0.07 | 1.119 ± 0.054 | 1.120 ± 0.044 | 0.988 |
